# Seeing Nemo: molecular evolution of ultraviolet visual opsins and spectral tuning of photoreceptors in anemonefishes (Amphiprioninae)

**DOI:** 10.1101/2020.06.09.139766

**Authors:** Laurie J. Mitchell, Karen L. Cheney, Wen-Sung Chung, N. Justin Marshall, Kyle Michie, Fabio Cortesi

## Abstract

Many animals can see ultraviolet (UV) light (shorter than 400 nm) undetectable to human vision. UV vision may have functional importance in many taxa including for foraging and communication in birds, reptiles, insects and teleost fishes. Shallow coral reefs transmit a broad spectrum of light and are rich in UV; driving the evolution of diverse spectral sensitivities in teleost reef fishes, including UV-sensitivity. However, the identities and sites of the specific visual genes that underly vision in reef fishes remain elusive and are useful in determining how molecular evolution has tuned vision to meet the ecological demands of life on the reef. We investigated the visual systems of eleven anemonefish (Amphiprioninae) species, specifically probing for the molecular pathways that facilitate UV-sensitivity. Searching the genomes of anemonefishes, we identified a total of seven functional visual genes from all five vertebrate opsin gene subfamilies. We found rare instances of UV-sensitive *SWS1* opsin gene duplications, that produced two functional paralogs (*SWS1α* and *SWS1β*) and a pseudogene. We also found separate *RH2A* opsin gene duplicates not yet reported in the family Pomacentridae. Finally, we report on both qualitative and quantitative aspects of opsin gene expression found in the adult retina of the false clown anemonefish (*Amphiprion ocellaris*), and their photoreceptor spectral sensitivities measured using microspectrophotometry.

## INTRODUCTION

Ultraviolet (UV) vision is widespread across the animal kingdom (Jacobs, 1992; Tovée, 1995), and is relied upon for many essential behaviours including foraging (Church et al. 1998; Siitari et al. 2002; Boulcott & Braithwaite, 2004; Novales Flamarique, 2013), mate selection (Bennett et al. 1997; Andersson & Amundsen, 1997; Smith et al. 2002; Rick et al. 2007) and detecting potential competitors (Rick & Bakker, 2008; Siebeck et al. 2010; Bohórquez-Alonso et al. 2018). Underlying vision are the opsins: G-protein-coupled receptors that bind via a Schiff base linkage to a carotenoid-derived (Vitamin A1 or A2) chromophore to form visual pigments mediating light absorbance from around 300-700nm (Collin et al. 2009). UV-sensitivity is mediated by visual pigments with peak sensitivities shorter than 400 nm and are found in many vertebrates (Bowmaker, 2008), including multiple teleost lineages (Lin et al. 2017). However, for many teleost fishes we still lack detailed information on the identities, sites and molecular evolution of specific genes and regulatory pathways that facilitate vision, including UV-sensitivity.

Visual opsins from all five vertebrate opsin subfamilies can be found in teleosts, including the UV-sensitive or very-short-wavelength-sensitive 1 (SWS1), short-wavelength-sensitive 2 (SWS2), medium-wavelength-sensitive (MWS), rhodopsin 1 (rod opsin, RH1) and rhodopsin-like 2 (RH2), and long-wavelength-sensitive (LWS) opsins (Bowmaker, 1990; Bowmaker, 2008). All of these families arose from an ancient vertebrate opsin that underwent multiple whole-genome and individual gene duplication events (Van de Peer et al. 2009), the latter of which facilitated the acquisition of novel visual opsins in multiple teleost lineages (Hofmann & Carleton, 2009; Rennison et al. 2012; Musilova et al. 2019). Further changes to opsin gene function can include their preservation and/or resurrection via gene conversion, a homogeniser and repair mechanism of paralogous genes (Katju & Bergthorsson, 2010; Cortesi et al. 2016). Currently most of what is known on teleost opsin gene duplications pertains to the *SWS2, RH1*, *RH2* and *LWS* subfamilies for which functional paralogs are found in many species (Rennison et al. 2012; Cortesi et al. 2016), whereas the duplication and retention of *SWS1* paralogs is considerably rarer (Rennison et al. 2012; Lin et al. 2017; Musilova et al. 2019).

UV-sensitivity in the teleost retina is conveyed by the expression of *SWS1* and/or *SWS2* opsin in the outer-segments of single cone cells, while sensitivity to longer-wavelengths is facilitated by the expression of *RH2* and/or *LWS* in double cone cells (i.e. two fused single cones) (Carleton et al. 2005, 2008; Parry et al. 2005; Spady et al. 2006; Dalton et al. 2015; Härer et al. 2018; Stieb et al. 2019). Although a switch from an A1 to A2 chromophore mediated by the cytochrome P450 enzyme, Cyp27c1 (Enright et al. 2015) can induce a substantial (~60 nm) shift in the peak spectral absorbance (λ_max_) of a visual pigment (Govardovskii et al. 2000), it is changes in the opsin protein that drive most of the variation in spectral tuning (Nickle & Robinson, 2007; Carleton et al. 2020), particularly in the UV-region (Yokoyama et al. 2016). More specifically, the interaction between the opsin protein and lysine residues (Lys296) on the surfaces of the chromophore binding-pocket determine the visual pigment’s wavelength of maximum absorption sensitivity (Yokoyama, 2000). Changes in the polarity and/or charge of amino acid residues at ‘key-tuning’ sites near the binding pocket can induce a short- or long-wavelength shift in spectral sensitivity (Carleton et al. 2005; Terai et al. 2006; Carleton, 2009). The cumulative tuning effects of these sites may be used to infer the peak spectral absorbance of visual pigments (Yokoyama et al. 2008), including SWS1-based pigments (Shi & Yokoyama, 2003; Yokoyama et al. 2016). Further alterations to the spectral tuning of vision can be achieved by differential opsin expression (Fuller et al. 2004; Johnson et al. 2013; Cortesi et al. 2016; Shimmura et al. 2017) and/or coexpression of opsins in the retina (Cheng & Flamarique, 2007; Dalton et al. 2014, 2017; Cortesi et al. 2016; Lührmann et al. 2019; Stieb et al. 2019). The combined effect of these visual tuning mechanisms on photoreceptor spectral absorbance are measurable using microspectrophotometry (MSP); a technique that combines microscopy and spectrophotometry (Dartnall et al. 1983).

Many small bodied teleosts that inhabit shallow environments, rich in UV-wavelengths such as coral reefs often possess *SWS1* opsin genes contained in single cones (Stieb et al. 2016, 2017; Lin et al. 2017; Musilova et al. 2019), and have UV-transmissive lenses (Losey et al. 2003; Marshall et al. 2006). UV-vision in reef fishes may aid the detection of UV-reflecting zooplankton prey (Stieb et al. 2017) and/or facilitate a short-distance communication channel hidden from predators, most of which lack UV-sensitive photoreceptors (Losey et al. 1999; Siebeck, 2004; Marshall et al. 2006; Siebeck et al. 2010). However, despite the widespread nature of UV-sensitivity and its importance in reef fishes, its genetic basis remains largely uncharacterised, and therefore was the aim of this study.

To do this, we used anemonefishes (subfamily, Amphiprioninae), which are an iconic group of reef fishes that obligately associate with one or more species of sea anemone. They are also sequential hermaphrodites living in strict social hierarchies governed by body size (Buston, 2003). The visual system of *Amphiprion akindynos* has been previously characterised revealing UV-sensitive single cones (~400 nm lambda max/λ_max_) containing SWS1 opsin, in addition to other spectral sensitivities (520, 498, 541 nm λ_max_) (Stieb et al. 2019).

The public availability of short- and long-read sequenced genomes for eleven species of anemonefishes has made it possible to study in detail the evolution of *SWS1* opsin genes in this group of reef fishes. We first identified the *SWS1* opsin genes and other visual opsin genes found in the genomes of eleven anemonefish species and provide information from their synteny analysis and phylogenetic reconstruction. Second, we identified the visual transduction and shut-off pathway genes that regulate opsin activity to show their synteny in all examined species. Third, we measured the opsin gene expression levels and photoreceptor spectral sensitivities in the retina (using MSP) of the false-clown anemonefish (*Amphiprion ocellaris*). This fundamental information on anemonefish vision will enable whole-organism investigations relating visual genes with the retinal neuroanatomy and behavioural ecology of a reef fish.

## MATERIAL AND METHODS

### Anemonefish visual opsin genes: identification, phylogeny and synteny

All genetic sequence analyses including the visual inspection, mapping and alignment of genes were performed using Geneious Prime (v. 2019.2.1). Our *in-silico* searches of anemonefish visual opsin genes involved annotating the regions containing the *SWS1*, *SWS2*, *RH2*, *LWS* and *RH1*opsin genes, along with their immediate upstream and downstream (flanking) genes in the genomes of eleven species of anemonefish. The publicly available assembled genomes for anemonefishes and a sister species, the lemon damselfish, *Pomacentrus moluccensis*, were accessed from various sources including for *A. ocellaris* (Tan et al. 2018; accession no. NXFZ00000000.1), *A. percula* (Lehmann et al. 2018; “Nemo Genome DB, 2018”; accession no. GCA_003047355.1), *A. frenatus* (Marcionetti et al. 2018; https://doi.org/10.5061/dryad.nv1sv), *A. akallopisos*, *A. melanopus*, *A. perideraion*, *A. nigripes*, *A. bicinctus*, *A. polymnus*, *A. sebae* and *Premnas biaculeatus* (Marcionetti et al. 2019). Reference gene sequences from *O. niloticus* were used to detect anemonefish visual genes including opsin genes (accession no. AY775108; AF247128; JF262086; JF262087), visual transduction pathway genes and shutoff genes (see Tab. S1-3) by mapping individual exons using low specificity (<30% similarity) against anemonefish genomes. Annotated gene sequences were confirmed by BLAST search in ENSEMBL v. 97 (accession date: 15.08.2019) (Zerbino et al. 2018) against the genomes of *D. rerio* (assembly no. GRCz11), *G. aculeatus* (assembly no. GCA_006229165.1) and *O. niloticus* (assembly no. GCA_001858045.3). Visual transduction and shut-off pathway genes were also identified in the *A. percula* and *A. ocellaris* genomes using predicted gene sequences obtained from Ensembl (ensembl.org) and their coding sequences were confirmed by assembly against the transcriptome of the *A. ocellaris* retina.

Phylogenetic trees based on the nucleotide alignment (MAFFT v. 7.388; Katoh & Standley, 2013) of 149 visual opsin coding sequences were generated using Bayesian inference in MrBayes v.3.2.7a (Huelsenbeck & Ronquist, 2001) in a workflow run through CIPRES (Miller et al. 2010) and viewed using Figtree v. 1.4.4 (Rambaut, 2018) that generated node support values. We included additional opsin sequences from unrelated species to show the grouping of anemonefish opsin genes relative to those found in other vertebrates obtained from genbank (www.ncbi.nlm.nih.gov/genbank/) including *Anolis carolinensis* (accession no. NM_001293118), *Oreochromis niloticus* (AY775108, JF262088, JF262086, JF262087), *Pseudochromis fuscus* (accession no. KP004335, KP017247), *Danio rerio* (AB087811, HM367062, AB087803, NM_001002443, AF109369, KT008394, NM_182892, NM_131254, KT008398, KT008399), *Oryzias latipes* (AB180742, XM_004069094, NM_001104694, AB223056, AB223057) and *Gasterosteus aculeatus* (KC774627, KC774623, KC594701, KC774625, KC774626). Two opsin trees were reconstructed: *1*) overall opsin tree in a GTR+I+G model selected based on the best-fit model AIC from Jmodeltest2 (Darriba et al. 2012; see supplementary materials for output) with default parameters, and *2*) an identical model, but with *SWS1* sequences restricted to only the fourth exon and intronic region between the fourth and fifth exons; where the majority of sequence variation exists between the two coding *SWS1* genes. Both trees used *Anolis carolinensis* vertebrate ancestral (VA) opsin as an outgroup. The Bayesian reconstructions included MCMC searches for 10 million generations with two independent runs and four chains each, a sampling frequency of 1000 generations and a burn-in of 25%.

### Anemonefish opsin gene conversion analysis

Visual opsin gene duplicates were tested for gene conversion: a phenomenon commonly found in teleosts (Owens et al. 2009; Watson et al. 2010; Nakamura et al. 2013; Cortesi et al. 2015; Sandkam et al. 2017; Escobar-Camacho et al. 2017, 2020). This was analysed using the program GARD (Genetic Algorithm Recombination Detection) (Kosakovsky Pond et al. 2006) on the aligned whole sequences of both the *RH2A* and *SWS1* duplicates in *A. ocellaris* and *A. percula* to reveal the presence/absence of recombination. Identified sites of recombination were corroborated using different phylogenetic tree topologies based on whole opsin gene sequences, and sequences limited to suspected regions of recombination.

### Animals and ethics statement

We used 13 captive-bred *A. ocellaris* (supplier Gallery Aquatica, Wynnum, 4178 QLD, Australia) for visual gene expression analysis, lens transmission measurements and microspectrophotometry (described below). Anemonefish were housed in recirculating aquaria at the Queensland Brain Institute at The University of Queensland, Australia. Experiments were conducted in accordance with The University of Queensland’s Animal Ethics Committee guidelines under the approval numbers: QBI/304/16 and SBS/077/17.

### Visual gene expression analysis of A. ocellaris

Adult retinae from two female (mean standard-length = 45 mm) and two male (mean standard-length = 30 mm) *A. ocellaris* were sampled for opsin gene expression analysis. All fish were kept under standard aquarium lighting for a minimum of three weeks (Fig. S1). Isolated retinas were homogenized using a high-speed bench-top homogenizer and total RNA was extracted using the QiaGen RNeasy Mini Kit. RNA was purified from any possible DNA contamination by treating samples with DNase following the protocol outlined by the manufacturer (QiaGen). The integrity of the extract was subsequently determined using a Eukaryotic Total RNA Nanochip on an Agilent 2100 Bioanalyzer (Agilent Technologies). Total RNA was sent to Novogene (https://en.novogene.com/) for library preparation and strand-specific transcriptome sequencing on a Hiseq2500 (PE150, 250~300 bp insert). Retinal transcriptomes were then filtered and transcripts *de novo* assembled on a customised Galaxy (v2.4.0.2; usegalaxy.org) (Afgan et al. 2016) workflow following the protocol described in de Busserolles et al. (2017). Individual cone opsin expression levels for single and double cones were then calculated separately, as per de Busserolles *et al.* (2017), where the number of mapped reads for each opsin gene was normalised by the total number of reads mapped and divided by its length. Rod versus cone opsin expression was calculated as the total proportion out of all mapped opsin reads.

To determine whether the identity of the chromophore could have a role in the tuning of MWS/LWS cone spectral sensitivities, we calculated the gene expression levels of the protein-encoding gene *CYP27C1*; a catalyst in the conversion of an A1 to A2 chromophore (Enright et al. 2015). We searched and annotated *A. ocellaris CYP27C1* by mapping *D. rerio CYP27C1* against the *A. ocellaris* genome. The coding regions of *CYP27C1* were mapped against retinal transcriptome assemblies and relative expression was quantified as the number of mapped reads per one million reads, as per Härer et al. 2018.

### Lens transmission and photoreceptor spectral sensitivities of *A. ocellaris*

For the measurement of lens transmission in *A. ocellaris*, the lenses (n=3 fish) were isolated and rinsed in PBS to remove any blood and vitreous. Spectral transmission (300-800 nm) was then measured by mounting the lens on a drilled metal plate between two fibres connected to an Ocean Optics USB2000 spectrometer and a pulsed PX2 xenon light source (Ocean Optics). Light spectra were normalised to the transmission value at 700 nm (Siebeck & Marshall, 2001), and lens transmission values were taken at the wavelength at which 50% of the maximal transmittance (T_50_) was attained (Douglas & McGuigan, 1989; Siebeck & Marshall, 2001).

The spectral absorbance of *A. ocellaris* photoreceptors were measured in a total of nine fish using single-beam wavelength scanning microspectrophotometry (MSP). This procedure followed that outlined elsewhere (Levine & MacNichol, 1979; Cheney et al. 2009; Chung & Marshall, 2016). In short, small pieces (~1 mm^2^) of tissue were excised from the eyes of two-hour dark-adapted fish, then immersed in a drop of 6% sucrose (1X) PBS solution and viewed on a cover slide (sealed with a coverslip) under a dissection microscope fitted with an infra-red (IR) image converter. A dark scan was first taken to control for inherent dark noise of the machine and a baseline scan measured light transmission in a vacant space free of retinal tissue. Pre-bleach absorbance measurements were then taken by aligning the outer segment of a photoreceptor with the path of an IR measuring beam that scanned light transmittance over a wavelength range of 300-800 nm. Photoreceptors were then bleached using bright white light for one minute. Successful (pre-bleach) cell scans were then re-measured and compared to photo-bleached scans of the same cell to confirm the observed light absorption was by a visual pigment. Confirmed visual pigment spectral absorbance data were then fitted in a custom (Microsoft Excel) spreadsheet using bovine rhodopsin as a template pigment to estimate the maximum absorbance (λ_max_) value of the visual pigment (Govardovskii et al. 2000; Partridge & De Grip, 1991). The quality of fit of absorbance spectra between A1- and A2-based visual pigment templates were also visually compared. Individual scans were binned based on their grouping of similar (≤10 nm difference) λ_max_ values and then averaged across individuals.

Comparisons between anemonefish opsin protein sequences and those of other fishes with known λ_max_ values were made to infer the spectral tuning effects of individual amino acid sites (see supplementary material). Estimates of anemonefish opsin λ_max_ values were calculated from the known spectral absorbances of *O. niloticus* (Parry et al. 2005), *Oryzias latipes* (Matsumoto et al. 2006), *Lucania goodei* (Yokoyama et al. 2007), *Maylandia zebra* (Spady et al. 2006), *Dascyllus trimaculatus* (Hofmann et al. 2012) and *Pomacentrus amboinensis* (Siebeck et al. 2010). These opsins have been thoroughly studied using either MSP or the *in-vitro* reconstitution of opsin proteins for a direct measurement of pure protein spectral absorbance (Carleton 2009; Matsumoto et al. 2006; Yokoyama & Jia, 2020). Our analysis involved identifying variable amino acid residues located at sites within the retinal binding pocket attributed to a shift in polarity and/or substitutions at previously reported tuning sites (Nakayama & Khorana, 1991; Fasick et al. 1999; Neitz et al. 1991; Janz & Farrens, 2001; Nagata et al. 2002; Shi & Yokoyama, 2003; Carleton et al. 2005; Parry et al. 2005; Spady et al. 2006; Yokoyama et al. 2007; Yokoyama, 2008; Yokoyama & Jia, 2020; Lührmann et al. 2019). Opsin protein sequences were aligned using MAFFT alignment (MAFFT v. 7.388; Katoh & Standley, 2013) with bovine (*Bos taurus*) rhodopsin as a template (PDB accession no. 1U19).

Visual pigment spectral absorbance curves were modelled and fitted against measured absorbance spectra for the (co)expression of opsins. The shape of the absorbance curve was calculated according to standard A1 and A2 chromophore templates (see Govardovskii et al. 2000) for each visual pigment based on estimated λ_max_ values. For modelling coexpression, the individual absorbance curves were multiplied by varying input proportions to simulate different amounts contained in the cone cells until the summed absorbance curve had a λ_max_ value close to that measured, while still fitting the shape of the cone absorbance spectra, with particularly close adherence in the long-wavelength limb.

## RESULTS

### Anemonefish visual opsin genes: identification, phylogeny and synteny

Our *in-silico* searches in the genomes of anemonefishes and one damselfish (*P. moluccensis*) yielded a total of eight fully-coding opsin genes belonging to five opsin classes including one rod opsin *RH1*, two ultraviolet-sensitive *SWS1* opsins, a single violet-sensitive *SWS2* opsin, three blue-green-sensitive *RH2* opsins, and a single yellow-red-sensitive *LWS* opsin (Fig. 1; Tab. S1). All visual transduction pathway genes and shutoff genes present in other vertebrates/fish species were also identified in *A. ocellaris* and *A. percula*, with no extra duplicates for these genes found (Tab. S2).

**Figure. 1.**
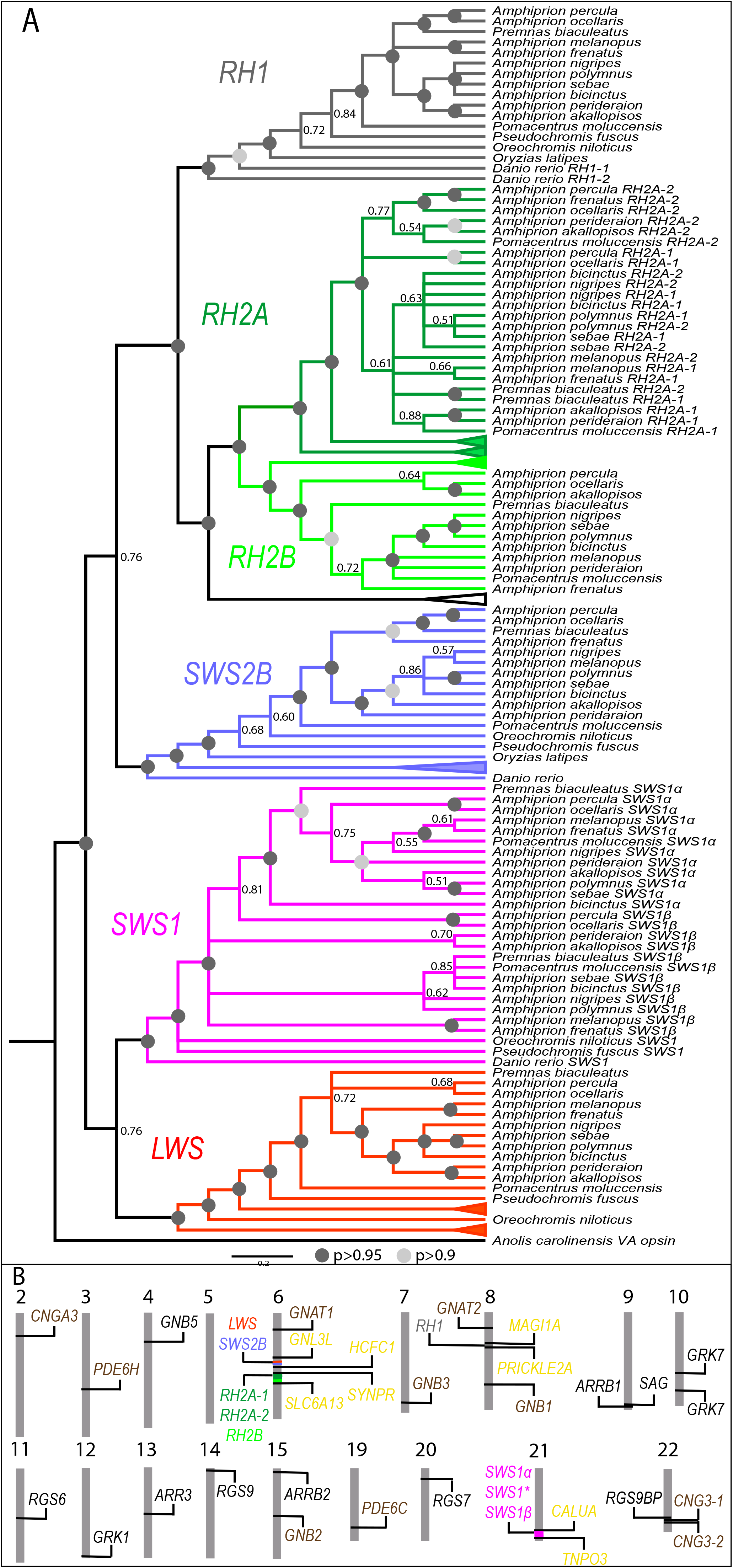
(A) Anemonefish visual opsin gene phylogeny reconstructed using the fourth exon and intron of *SWS1α* and *SWS1β*. Shown are internodal Bayesian posterior probabilities (P) depicted as ‘dark grey’ and ‘light grey’ markers that indicate posterior probabilities greater than 0.95 and 0.9, respectively. Posterior probabilities less than 0.9 are given in full. For the purpose of presentation, all clustered outgroup branches were collapsed. To view the complete non-collapsed phylogeny, see the supplementary material. (B) Synteny of anemonefish visual genes found in the genomes of *A. percula* and *A. ocellaris* including opsin genes and their flanking genes (yellow), visual transduction pathway genes (brown), visual cascade shutoff pathway genes (black) and their corresponding chromosome number. Opsin gene acronyms stand for RH1 = Rhodopsin 1 (rod opsin), RH2 = Rhodopsin 2, SWS2 = Short-wavelength-sensitive 2, LWS = Long-wavelength-sensitive, SWS1* = Short-wavelength-sensitive 1/Short-wavelength-sensitive pseudogene, va = vertebrate ancient opsin (outgroup). Visual transduction and shutoff pathway gene acronyms stand for CNG = Cyclic nucleotide gated channel, PDE6 = Phosphodiesterase 6, GNAT = G protein subunit alpha transducin, GNB = G protein subunit beta, ARR = Arrestin, ARRB = Arrestin beta, SAG = S-antigen visual arrestin, GRK = G protein-coupled receptor kinase, RGS = Regulator of G-protein signalling.

Two UV-sensitive opsins, *SWS1α* and *SWS1β* were identified in all examined anemonefishes. Those found in *A. percula* and *A. ocellaris*, matched the sequence predictions on ensembl (ensembl transcript id: ENSAOCT00000030401.1, ENSAOCT00000007343.1, ENSAPET00000035041.1, ENSAPET00000035052.1). Protein sequence analysis revealed highly conserved SWS1 opsins, where both encoded 339 aa (Fig. S2) and shared 96% similarity. Closer inspection of the protein sequences showed that 15% of the differences between paralogs occurred at known SWS1 tuning sites (Yokoyama, 2008; Shi & Yokoyama, 2003), and 90% were within the seven transmembrane domains. Two of these sites were found to have a nonpolar to polar shift including A118S and A114S, with the former known to induce a +5 nm shift in opsin λ_max_ (Shi & Yokoyama, 2003).

All SWS1 protein sequences had conserved F86 and S90 aa sites, an invariable feature of UV-sensitive opsins (Cowing et al. 2002). The *A. percula* genome showed that both *SWS1* genes were located in-tandem on Chromosome 21, separated by 19kbp and spanning a total syntenic region of 29kbp. A third non-functional *SWS1* pseudogene (Fig. 1B) was also found between the two paralogs (encoding 196 amino acids), with highest sequence homology to the initial two exons of *SWS1β*. In both species, the *SWS1* pseudogenes were found to have an 81 base pair insertion near the 3’ end of Exon 1 that contained a premature stop codon at aa number 123. The *SWS1* pseudogene in *A. ocellaris* was also found to have a missense mutation caused by a single Guanine to a Cytosine substitution that removed the start codon. The genes flanking the *SWS1* region were identical across all species.

All anemonefishes were found to have three blue-green sensitive opsin genes; *RH2A-1*, *RH2A-2* and *RH2B*, that closely matched sequence predictions on ENSEMBL (ensembl transcript id: ENSAOCT00000002771.1, ENSAPET00000018561.1, ENSAOCT00000002796.1, ENSAPET00000018598.1, ENSAOCT00000002830.1, ENSAPET00000018632.1). Like in other examined teleosts, the *RH2* genes (Fig. 1B) in *A. ocellaris* and *A. percula* were found in-tandem, spanning a region of approximately 29kbp and flanked by identical genes immediately up- and down-stream of the syntenic region. The translated protein sequences of both RH2A and RH2B (Fig. S2) encoded 353- and 346-amino acids (aa), respectively. *RH2A* paralogs were found to be highly conserved within *A. percula* and *A. ocellaris*, sharing 95.2% (*RH2A-1*) and 98% (*RH2A-2*) similarity, respectively.

The chromosomal resolution of the *A. percula* genome revealed the locations of the *RH2* and *LWS*-*SWS2B* syntenic regions (Fig. 1B) in-tandem on Chromosome 6, separated by approximately 8.6mbp; a conserved syntenic region shared by many other neo-teleost species (Musilova et al. 2019). One red-sensitive *LWS* opsin and violet-sensitive, *SWS2B* opsin was identified in both *A. percula* and *A. ocellaris* that matched sequence predictions on ensembl (ensembl transcript id: ENSAOCT00000024298.1, ENSAPET00000033644.1, ENSAOCT00000031935.1, ENSAPET00000033670.1). The translated protein sequences of LWS and SWS2B were found to encode 358- and 352-aa, respectively.

Phylogenetic analyses placed all the identified opsin genes into distinct homologous clusters with their predicted opsin classes (Fig. 1A). Furthermore, the translated and aligned protein sequences for the identified genes (Fig. S2) exhibited typical opsin traits including the conserved chromophore binding site residue (K296) and in most cases intact seven transmembrane domains; however, the latter was only found for all opsin genes in the *A. ocellaris* and *A. percula* genomes, whereas in other species only partial (incomplete) gene sequences for some opsins could be found including for *RH2A-2* and *SWS1α* opsin genes. It was for this reason, in addition to the presence of premature stop codons in most species’ *SWS1β* opsin gene that translated protein sequences of *RH2A-2* and *SWS1* genes were only examined in *A. percula*, *A. ocellaris* and (*SWS1β* in) *A. frenatus*. Whether these incomplete/pseudogenized genes are a true biological occurrence or sequencing errors as a result of short-read assembled draft genomes remains to be investigated. Initial *SWS1* phylogeny based on complete coding sequences revealed a partial clustering of two gene groups; the paralogs *SWS1α* and *SWS1β* (Fig. S3). Re-construction of the phylogeny used only Exon 4 and part of the intron between Exons 4 and 5, where most of the informative sites (i.e. single nucleotide polymorphisms, ‘SNPs’) were located, and provided a clearer separation of the paralogs into two distinct clusters (Fig. 1A). Phylogenetic analysis of *SWS2B*, *RH2B* and *LWS* in anemonefishes showed a typical pattern of species-relatedness that mostly resembled the inferred phylogenies reported elsewhere (Litsios et al. 2014; Rolland et al. 2018). Like most teleost fishes (Musilova *et al*. 2019), anemonefishes were found to possess a single *RH1*, with a conserved *RH1* syntenic region. The *RH1* phylogeny closely adhered to the species phylogeny (Fig. 1A).

### Anemonefish opsin gene conversion analysis

We found evidence of gene conversion in both the *RH2A* and *SWS1* duplicates in *A. ocellaris* and *A. percula* (Fig. 2). GARD analysis revealed two major breakpoints in *RH2A-1* and *RH2A-2* located at 634-892 bp (exons 3 and 4) and 893-1,059 bp (exons 4 and 5), that indicated recombination had occurred within the last few exons. Alternative tree topologies based on different *RH2A* breakpoint regions (Fig. 2) supported one of the two regions (892-1,059 bp) having recombination, as evident by the orthologous grouping of *RH2A* genes rather than paralogous grouping that was observed when tree topologies were based on either the 634-892 bp region or complete coding sequence. Analysis of the aligned RH2A opsin protein sequences identified only one known key tuning site (aa site 292; site number given according to bovine rhodopsin) (Yokoyama & Jia, 2020) located within the region of recombination.

**Figure. 2.**
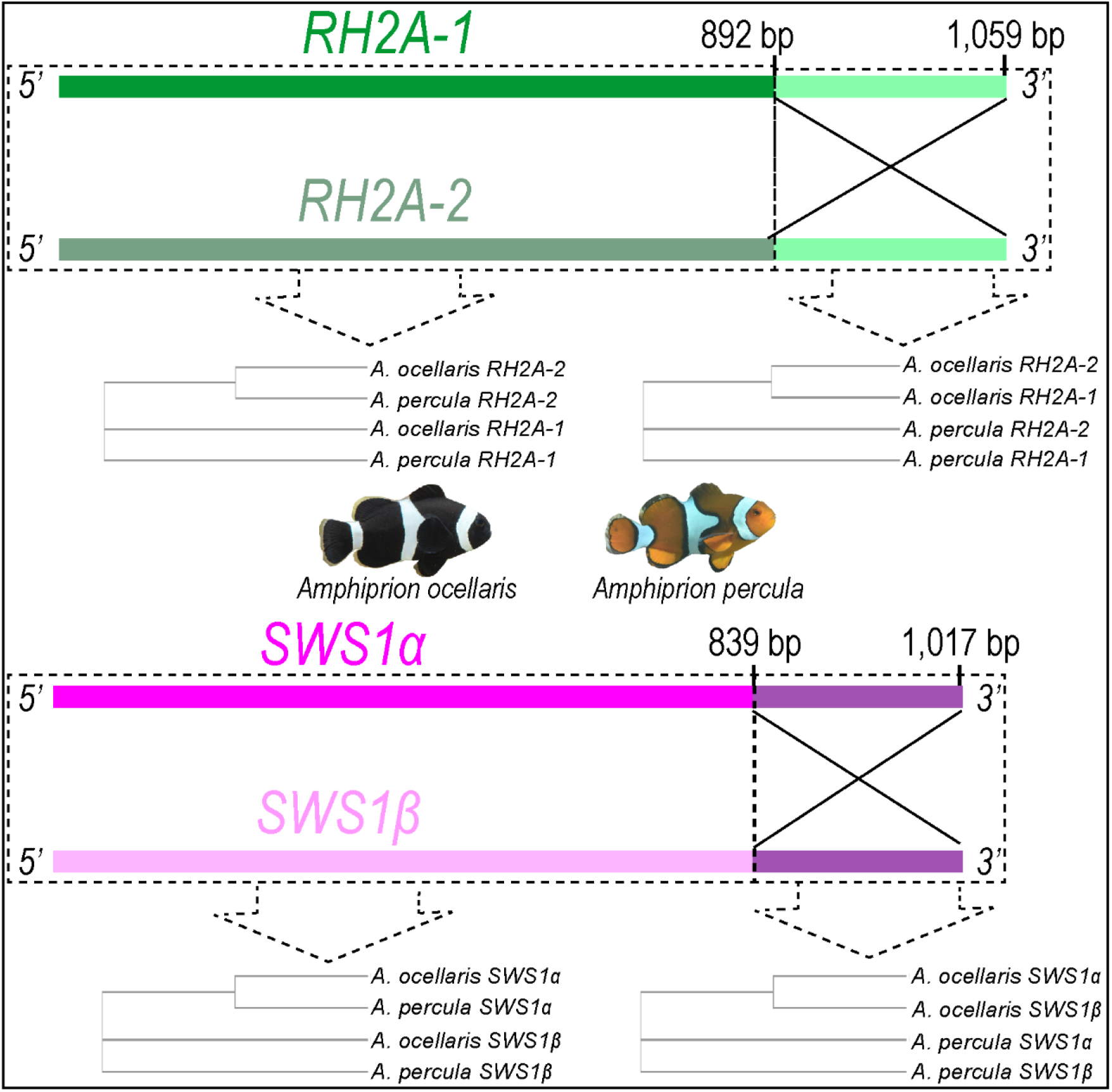
Regions of gene conversion detected in anemonefish (*A. percula* and *A. ocellris*) RH2A (*RH2A – 1* and *RH2A – 2* duplicates and SWS1 (*SWS1α* and *SWS1β*) duplicates. Boxes outline the alternative tree topologies constructed when using either the entire coding sequence or only the region of gene conversion.

One major breakpoint was found in *SWS1α* and *SWS1β* indicating that recombination occurred in the last two exons (839-1,017 bp). Alternative tree topologies supported the notion of gene conversion within this breakpoint region (Fig. 2), where opsin copies grouped as orthologs or as paralogs depending on whether trees were constructed using the breakpoint region or the complete coding sequence, respectively. Analysis of the aligned SWS1 opsin protein sequences did not yield any known tuning site changes within the recombined region.

### Visual gene expression analysis of *A. ocellaris*

Analyses of retinal transcriptomes (Fig. 3) revealed that under aquarium lighting the *A. ocellaris* retina (n=4) expressed one rod opsin; *RH1* (mean + s.d. = 70.63 ± 11.51%), and six cone opsins including four double cone opsins; *RH2A-1* (43.13 ± 6.68%.), *RH2B* (35.34 ± 3.21% *LWS* (11.45 ± 6.77%) and *RH2A-2* (10.08 ± 11.66%), and two single cone opsins; *SWS1β* (59.09 ± 9.41%.) and *SWS2B* (40.8 ± 8.70%) (Fig. 3). No trace of *SWS1α* expression was detected in three out of the four retinae and only a low amount (1.5%) was detected in one retina. No difference in opsin expression was found between males (n=2) and females (n=2) (Pairwise ANOVA, *p*-value > 0.05).

**Figure. 3.**
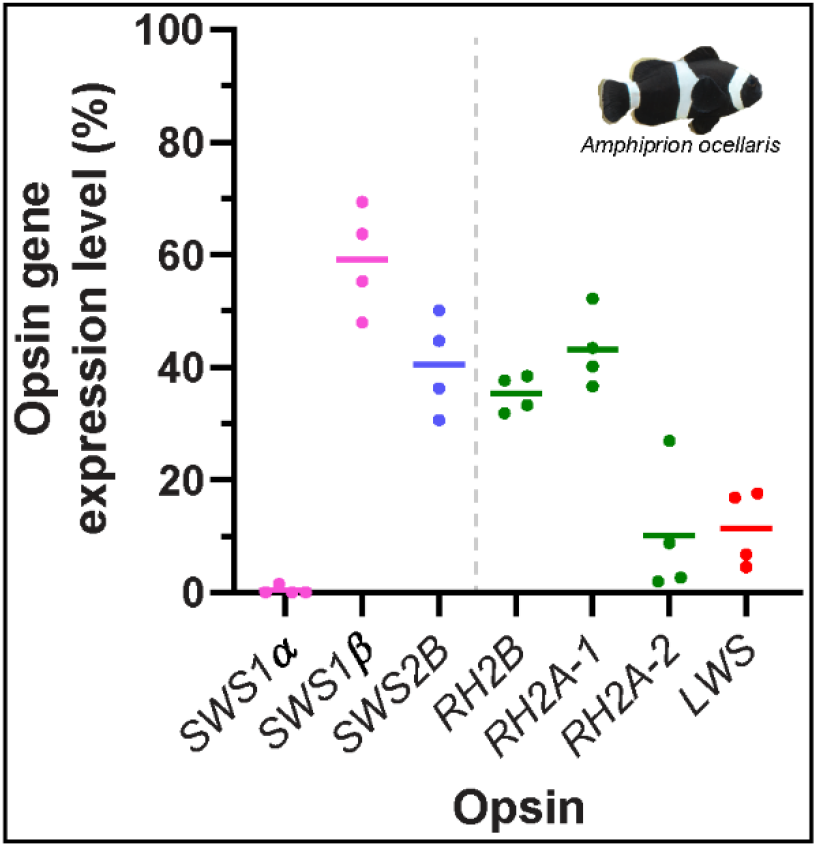
Relative cone opsin expression levels in the retina of captive *A. ocellaris* (N=4; 2 females and 2 males) kept under aquarium lighting (see supplementary material for illumination spectra). SWS1 = Short-wavelength-sensitive 1, SWS1 = Short-wavelength-sensitive 2, RH2 = Rhodopsin 2, LWS = Long-wavelength-sensitive. Lines represent the median proportion of opsin expression levels of single cones opsins (SWSs) and double cone opsins (RH2s, LWS), respectively. Individual points represent a smapled fish.

The gene *CYP27C1* was found to be expressed in very low amounts in all four *A. ocellaris* retinae, where relative expression levels ranged from 0.92 to 4.04 mapped reads per million reads.

### Lens transmission and photoreceptor spectral sensitivities of *A. ocellaris*

Transmission measurements on the lenses of *A. ocellaris* (individual T50 values = 320/339/362 nm) were averaged to give a mean T50 wavelength value of 341 nm (Fig. 4A). This T50 value was used to correct the photoreceptor spectral absorbance curves for the filtered light reaching the retina.

**Figure. 4.**
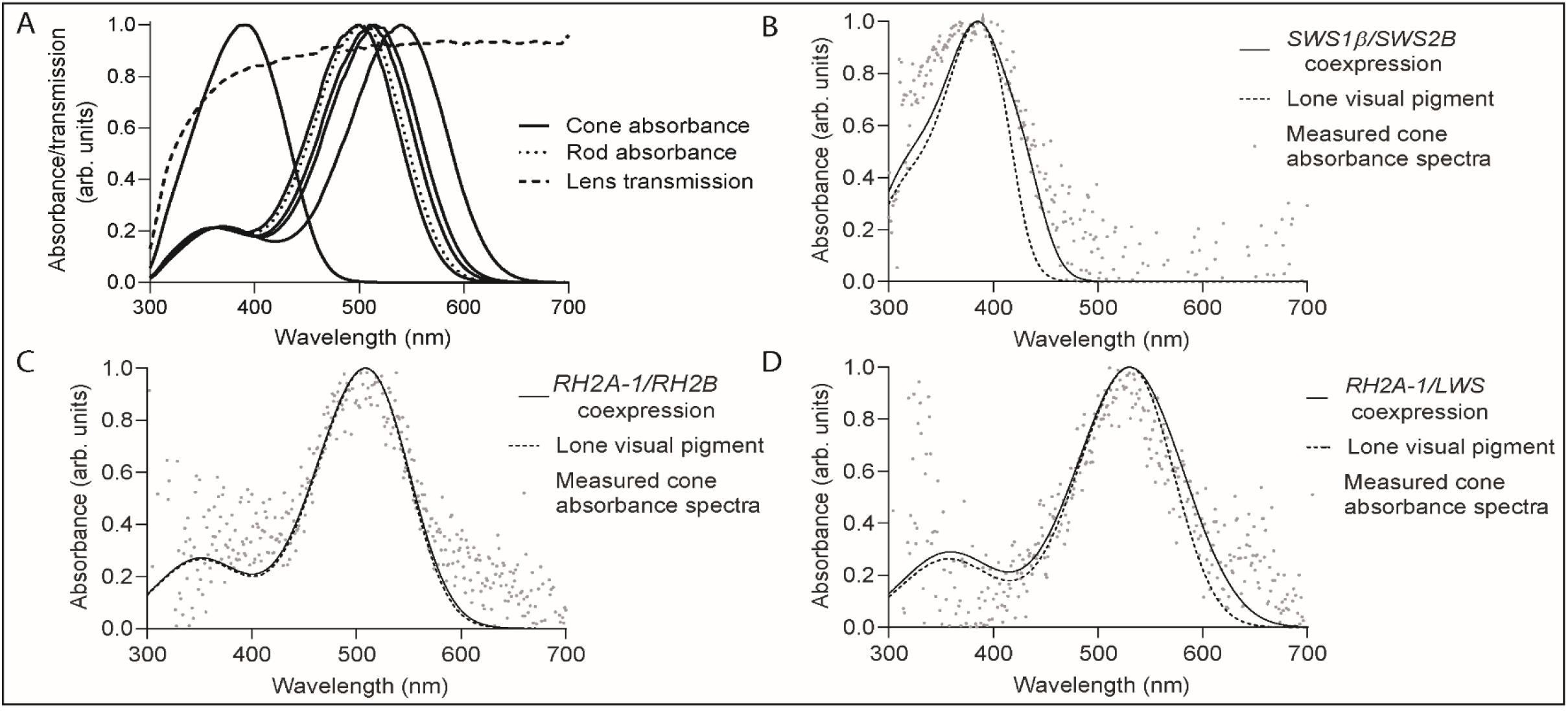
(A) Nomalised average lens transmission of A. ocellaris, and absorbance curves fitted to average rod and cone absorbance spectra measured in *A. ocellaris* (N=9). Modelled spectral absorbance curves for different cone opsin coexpression scenarios for (B) single cone (λmax = 385 nm) coexpressing SWS1α (40%) and SWS2α (60%), (C) double cone (λmax = 508.509 nm) coexpressing RH2B (35%) with RH2A-1 (65%), and (D) double cone (λmax = 531 nm) coexpressing RH2A-1 (55%) with LWS (45%). Plotted alongside for comparision are modelled spectral absorbance curves for lone visual pigments and actual measured cone absorbance spectra.

MSP in *A. ocellaris* (Fig. 4A) found one medium-wavelength-sensitive (MWS) rod (mean λ_max_ value = 502 ± 2.0 nm; n=10 cells, N = 4 fish; Fig. 4A), one short-wavelength-sensitive (SWS) single cone, three MWS double cones, and one LWS double cone. Otherwise mentioned, all measured absorbance spectra fitted an A1 visual pigment template.

Single cone λ_max_ measurements (mean λ_max_ value = 386 ± 4.6 nm; n=4, N = 4 fish; Fig. 4A) did not closely match any of the λ_max_ estimates for SWS1 (*SWS1α* estimated λ_max_ value = 356 nm; *SWS1β* estimated λ_max_ value = 370 nm; Fig. S2) or SWS2B (estimated λ_max_ value = 407 nm); however, modelling the absorbance curve from the coexpression of *SWS1β* (40%) and *SWS2B* (60%) provided a close fit (λ_max_ value = 384-386 nm; Fig. 4B). The single cone averaged absorbance spectra had an asymmetrical shape with a breadth more like an A2 visual pigment template.

One MWS double cone had λ_max_ measurements (mean λ_max_ value = 496.7 ± 0.58 nm; n=3, N = 3 fish; Fig. 4A) that closely matched the estimate for RH2B (estimated λ_max_ value = 497 nm; Fig. S2), while another double cone had λ_max_ measurements (mean λ_max_ value = 515 ± 2.1 nm; n=6, N = 3 fish; Fig. 4A) that closely matched one of the estimates for *RH2A-1/RH2A-2* (estimated λ_max_ value = 516 nm; Fig. S2). A separate λ_max_ for *RH2A-2* opsin could not be estimated, as the variable amino acid sites between the paralogs had unknown effects on spectral tuning and were absent in all reference sequences. One of these unknown variable sites, T266V exhibited a polarity shift that consistently alternated between the *RH2A* paralogs of *A. percula* and *A. ocellaris*.

Two measurements in one individual hinted at the presence of an intermediate MWS double cone (508/509 nm λ_max_, N = 2 fish; Fig. 4A), with a peak spectral sensitivity that did not match any visual pigment estimates. Although it was reproducible by modelling the coexpression of *RH2B* (35%) and *RH2A-1* (65%) (λ_max_ value = 507-509 nm; Fig. 4C), there was little difference to the modelled absorbance spectra of a lone visual pigment (Fig. 4C).

Similarly, the longer-wavelength sensitive double cone measurements (531/538 nm λ_max_, N = 2 fish; Fig. 4A) provided a poor match to the LWS opsin estimate (estimated λ_max_ value = 560 nm; Fig. S2) and could only be produced by modelling the coexpression of *RH2A-1* (55%) with *LWS* (45%) (λ_max_ value = 529-531 nm; Fig. 4D), and *RH2A-1* (40%) with *LWS* (60%) (λ_max_ value = 537-539 nm; Fig. S4), respectively. However, the latter provided an overall poor fit owing to the high amount of noise in the absorbance data that combined with only having two measurements made it also difficult to compare the fit between A1 and A2 visual pigment templates.

## DISCUSSION

Anemonefish genomes revealed visual opsin orthologs belonging to all five ancestral vertebrate classes of opsin (Davies et al. 2012) including *SWS1*, *SWS2*, *RH1*, *RH2*, and *LWS*, along with additional *SWS1* and *RH2A* duplicates. We found rare instances of UV-sensitive *SWS1* opsin gene duplication events that produced two functional paralogs (*SWS1α* and *SWS1β*) and a pseudogene. Moreover, the expressed cone opsin palette in the adult retinae of *A. ocellaris* comprised of *SWS1β* and *SWS2B* single cone opsins, in addition to *RH2A-1*, *RH2A-2*, *RH2B*, and a small amount of *LWS* in double cone opsin genes. Lens transmission measurements and MSP strongly implicate UV-sensitivity in *A. ocellaris* including the presence of UV-transmissive lenses and retinas that contain a single cone with a λ_max_ value of ~386 nm. Additional spectral sensitivities found by MSP in *A. ocellaris* included one rod with a λ_max_ value of 502 nm and four double cone spectral sensitivities with λ_max_ values of 497 nm, 515 nm, ~535 nm, and a possible fifth cone with a λ_max_ value of ~508 nm. We assume that *A. ocellaris* SWS opsins are only expressed in single cones, while the longer-wavelength sensitive RH2 and LWS genes are strictly-expressed in the double cones, as has been found in the anemonefish, *A. akindynos* (Stieb et al. 2019) and in cichlid fishes (Carleton et al. 2005, 2008; Parry et al. 2005; Spady et al. 2006; Dalton et al. 2015; Härer et al. 2019).

Visual opsin genes found in the genomes of anemonefishes are situated in conserved regions that share a common synteny with other teleost fishes (Lin et al. 2017). It also appears that anemonefish *SWS1* and *RH2* paralogs emerged through tandem duplication, as is common for opsin paralogs in many fish species (Cortesi et al. 2015; Lin et al. 2017; Musilova et al. 2019). It is extremely rare to find an *SWS1* opsin gene duplicate in teleosts (Rennison et al. 2012; Lin et al. 2017; Musilova et al. 2019). Here, we show that *SWS1* opsin duplicates are conserved across eleven members of Amphiprioninae and in a species of damselfish (*P. moluccensis*), as similarly reported in *Chromis chromis*; a member of the Pomacentrid subfamily, Chrominae (Musilova et al. 2019). Conversely, only one *SWS1* opsin gene was reported in the damselfish, *Stegastes partitus* (Musilova et al. 2019), and hence not all Pomacentridae possess *SWS1* opsin duplicates. Thus, it remains unclear whether the gene duplication event that produced *SWS1α* and *SWS1β* was lineage- or subfamily-specific. More genomic and/or transcriptomic data is needed to resolve whether this *SWS1* gene duplication occurred at the base of Pomacentridae or multiple times independently in its radiated lineages.

Interestingly, we found evidence of a second gene duplication event that produced a third psuedogenised *SWS1* paralog in the genomes of *A. ocellaris* and *A. percula*. This *SWS1* pseudogene was more alike *SWS1β* than *SWS1α*, suggesting it was either a duplicate of the former or gene conversion had occurred. The genomes of these species were published by two different lab-groups (Tan et al. 2018; Lehmann et al. 2018) and assembled differently, making it unlikely that this pseudogene is an assembly artefact. Due to the basal placement of these sister-species in the anemonefish phylogeny (Santini & Polacco, 2006), it remains possible this second duplication event occurred relatively early in anemonefish evolution. Alternatively, this duplication-event was specific to the clade containing *Premnas* and the sister-species, or even limited to only the sister-species. This pseudogene was not detected in the other eight anemonefish genomes; however, this was likely owing to these draft genomes being comparatively poorly resolved, that similarly meant only the fourth and fifth exons of *SWS1α* could be identified in those species. This incomplete or incorrect assembly of highly similar genomic regions is an inherent problem of short-read draft genomes.

Both SWS1 opsin protein sequences in *A. ocellaris* and *A. percula* had a conserved alanine at site 86 and serine at site 90; crucial for UV-sensitivity (Shi & Yokoyama, 2003; Tada et al. 2009). Furthermore, both anemonefish *SWS1* duplicates have intact open reading frames but only *SWS1β* is expressed in the adult retina. This combined with a UV-transmissive lens and presence of a UV/violet sensitive single cone provide a strong indication of UV-vision mediated by SWS1 and SWS2B visual pigments in the single cones. Like other damselfishes, the examined anemonefishes all possess the violet sensitive SWS2B opsin (Hofmann et al. 2012; Stieb et al. 2016; Stieb et al. 2017). Moreover, no indication for the percomorph-specific *SWS2Aα* and *SWS2Aβ* was found, likely due to a double-gene loss in the Pomacentrid ancestor (Cortesi et al. 2015).

The large mismatch between spectral sensitivity measurements and estimates of *A. ocellaris* single cones and their opsins (SWS1 and SWS2B) suggest the measured spectral sensitivities are not produced by the expression of either opsin alone. Rather, it is more likely that *SWS1β* and *SWS2B* are coexpressed in the single cones to produce an intermediate absorbance curve; a phenomenon supported by fitting the absorbance spectra to the modelled equal coexpression of both opsins. Moreover, the asymmetrical and broad absorbance curve of the *A. ocellaris* single cone resembles that found in other species such as *A. akindynos*, where the single cones have a broad absorbance curve and λ_max_ value of 400 nm, despite SWS1 and SWS2B having estimated λ_max_ values of 370 nm and 407 nm, respectively (Stieb et al. 2019). Likewise, the coexpression of SWS visual pigments has been reported in the single cones of African cichlids (Dalton et al. 2017). Although not examined in this study, there can also be spatial patterns in the coexpression of SWS opsins across the retina, such as in *A. akindynos*, where coexpression is localised to a small, dorso-temporal (i.e. forward-looking) area of high acuity that may aid specific tasks (Stieb et al. 2019). Together, the close to equal ratio of *SWS1α* and *SWS2B* expression levels in the *A. ocellaris* retina, along with the failure to find a pure UV-sensitive and/or violet-sensitive cone strongly supports the notion of opsin coexpression in the single cones. Future in-situ mapping of cone opsin expression across the *A. ocellaris* retina will resolve the spatial patterns of opsin (co)expression.

The importance of UV-vision in anemonefishes is unclear but looking to other UV-sensitive teleosts could direct further investigation. Like the anemonefishes, the rainbow trout (*Oncorhynchus mykiss*) has retained two *SWS1* opsins that originated from a family-specific whole-genome duplication event, and their UV-sensitivity improves zooplanktivory efficacy by enhancing prey contrast (Flamarique & Hawryshyn, 1994; Flamarique, 2013). Similar foraging benefits conveyed by UV-sensitivity have been shown in perch (Loew et al. 1993), cichlids (Jordan et al. 2004), sticklebacks (Rick et al. 2012), and zebrafish (Flamarique, 2016). Indeed, in damselfishes higher *SWS1* expression correlates strongly with zooplanktivory (Stieb et al. 2017). Anemonefishes are life-long zooplanktivores (Fautin & Allen, 1997) that could similarly benefit from the enhanced UV contrast of prey in the water column.

Fish also use UV-signals for communicating with rivals and mates, as reported in the guppy (Smith et al. 2002), stickleback (Rick et al. 2007, 2008) and a damselfish (Siebeck et al. 2010). In *A. akindynos*, the coexpression of *SWS1* and *SWS2* is believed to increase the chromatic contrast of its skin colour patterns that may improve the detection of conspecifics (Stieb et al. 2019). The use of UV-signalling in communication is worth further investigation in anemonefishes, as they possess UV-reflective skin patterns (Marshall et al. 2006; Stieb et al. 2019) that could conceivably be used to communicate their dominance status to other members within a family group and/or convey the occupied-status of their hosted anemone to the members of rival groups from nearby anemones.

Interestingly, the advantages of UV-vision could be conceivably conveyed by a single UV-sensitive opsin, and the functional significance of possessing SWS1 duplicates remains unknown. Another teleost species found to have two functionally coding *SWS1* duplicates is the smelt, *Plecoglossus altivelis* and like *A. ocellaris* it only expresses one paralog (Minamoto & Shimizu, 2005). As similarly suggested in *P. altivelis*, it is possible that the second *SWS1* paralog is expressed by *A. ocellaris* during specific seasons such as winter months on the reef, when there is higher ambient UV and lower turbidity promoting the more efficient transmission of shorter wavelengths of light. Increased SWS1 opsin expression has been found in the damselfish, *P. nagasakiensis* during winter months and may be a visual tuning response for taking advantage of the higher UV (Stieb et al. 2016). Seasonal changes in opsin gene expression levels have been reported to alter colour perception in widespread taxa (see review by Shimmura et al. 2018).

Analysing opsin expression levels in the larval and/or early-juvenile anemonefish retina may also reveal whether the shorter-wavelength sensitive *SWS1α* is expressed during earlier life stages, and whether an ontogenetic shift from *SWS1α* to *SWS1β* occurs as a response to change(s) in light environment, particularly during the settlement stage when pelagic larvae return to the reef to seek a host anemone. Ontogenetic changes in cone opsin expression levels as a response to change in habitat have been reported in other reef fishes including the spotted unicornfish (*Naso brevirostris*) (Tettamanti et al. 2019) and dusky dottyback (*P. fuscus*) (Cortesi et al. 2016). Alternatively, the *SWS1α* opsin may be expressed in other tissues and/or organs, where animal opsins have been demonstrated to serve other sensory modalities including non-image forming vision, thermosensation, hearing (see review by Leung & Montell, 2017) and most recently in taste (Leung et al. 2020).

The anemonefish double cones expressed a greater variety of opsins including two *RH2A* paralogs (*RH2A-1* and *RH2A-2*), *RH2B* and *LWS*, almost all of which have been found in other pomacentrids (Hofmann et al. 2012; Stieb et al. 2016; Stieb et al. 2017). However, to our knowledge, this is the first report of an *RH2A* duplication in a pomacentrid. Similar *RH2A* duplications have been reported in most percomorphs (as reviewed by Musilova et al. 2019). It was previously difficult to separate RH2A duplicates in pomacentrids due to their high degree of similarity, coupled with the low-resolution genomes available. Anemonefish *RH2A* duplicates share a high level of similarity partly due to gene conversion detected in the fourth and fifth exons; a region that may be preserved to maintain the opsin-chromophore binding site (Lys296), as is likely the case in *SWS1*. However, unlike the region of gene conversion found in *SWS1*, that found in *RH2A* also encompasses a known tuning site, bovine rhodopsin site number 292, where a large shift of −11 nm occurs when Ala is substituted with Ser (Yokoyama & Jia, 2020). Thus, preservation of this site by gene conversion may also have importance in maintaining the spectral tuning of RH2A visual pigments.

Based on only the spectral sensitivities of the double cones, the spatial patterns of opsin expression in the anemonefish retina cannot be inferred. However, we speculate the most likely patterns of opsin expression in anemonefish double cones. In *A. ocellaris*, it appears that *RH2A-1* opsin (λ_max_ value = 516 nm) and *RH2B* opsin (λ_max_ value = 497 nm) are expressed in separate double cone members, as indicated by the close matches between measurements of cone spectral absorbance and estimates of visual pigment tuning. It is unclear whether the two visual pigments are also coexpressed in another double cone to produce an intermediate spectral sensitivity (λ_max_ value = 508/509 nm), as the possible expression of a short-wavelength shifted *RH2A-2* opsin cannot be out-ruled. Multiple variable amino acid sites were found between the *RH2A* duplicates that could not be accounted for when estimating tuning effects on the opsin. Moreover, many of these substitutions exhibit changes in polarity suggesting a likely effect on spectral tuning, for example site T266V that alternated between paralogs and may be a key tuning site in anemonefish RH2A tuning. In cichlids, the differential expression of duplicate *RH2A* opsins in the double cones is believed to improve blue-green sensitivity that may aid in viewing nuptial skin colours and/or colonising different depths (Weadick & Chang, 2012; Dalton et al. 2014). Moreover, the coexpression of *RH2A* paralogs with either *RH2B* or *LWS* opsins have also been shown to enhance luminance contrast (Dalton et al. 2014). Modelling the coexpression of anemonefish *LWS* and *RH2A-1/RH2A-2* opsins indicated it was likely responsible for producing the longer-wavelength sensitive double cones measured in the *A. ocellaris* retina. The in-situ mapping of cone opsin expression across the anemonefish retina is needed to resolve spatial patterns of opsin gene expression. Alternatively, it is feasible that a switch from an A1 to A2 chromophore could contribute towards the MWS/LWS tuning of the double cones; however, this seems unlikely when considering the very small amounts of *CYP27C1* expression detected in the *A. ocellaris* retina that were a couple orders of magnitude below levels typically detected in cichlids that express an A2 chromophore (Härer et al. 2018).

## CONCLUSIONS AND FUTURE DIRECTIONS

Here we have shown that anemonefishes possess eight visual opsin genes including duplications of the *SWS1* and *RH2* genes. Moreover, most of these opsins were found expressed in the adult retinae of *A. ocellaris*. Our reported visual opsin expression levels provide an initial glance at the opsin expression profile in the retina of captive *A. ocellaris*, and therefore, comparisons with wild anemonefish are required to rule out differences associated with lighting and/or seasonality. The presence of two functional *SWS1* opsins in all examined anemonefishes suggests an anemonefish specific *SWS1* gene duplication event, and a possibly clade-specific second duplication event that produced an *SWS1* pseudogene. Circumstantial evidence including the expression of *SWS1* opsin, a UV-transmissive lens and presence of UV/violet-sensitive cone cells in the *A. ocellaris* retina strongly implicate UV-vision. Finally, we hope that our characterisation of the anemonefish visual genes will encourage future experiments using them as a model for studying reef fish vision.

## Acknowledgements

We would like to thank Gillian Lawrence and the University of Queensland Biological Resources Aquatics Team for their support in maintaining aquaria. We also thank Prof. Karen L. Carleton (University of Maryland, USA), Dr. Sara Marie Stieb, and Dr. Daniel Escobar-Camacho for their insight, and constructive feedback on the initial manuscript.

