## supplementary for "Seeing Nemo: molecular evolution of ultraviolet visual opsins and spectral tuning of photoreceptors in anemonefishes (Amphiprioninae)"

**Supplementary materials**

*Supplementary Figure 1*. Absolute downwelling irradiance levels of fluorescence lighting in anemonefish aquaria measured using a spectrometer and fibre fitted with a cosine corrector in anemonefish aquaria produced by fluorescent lighting.

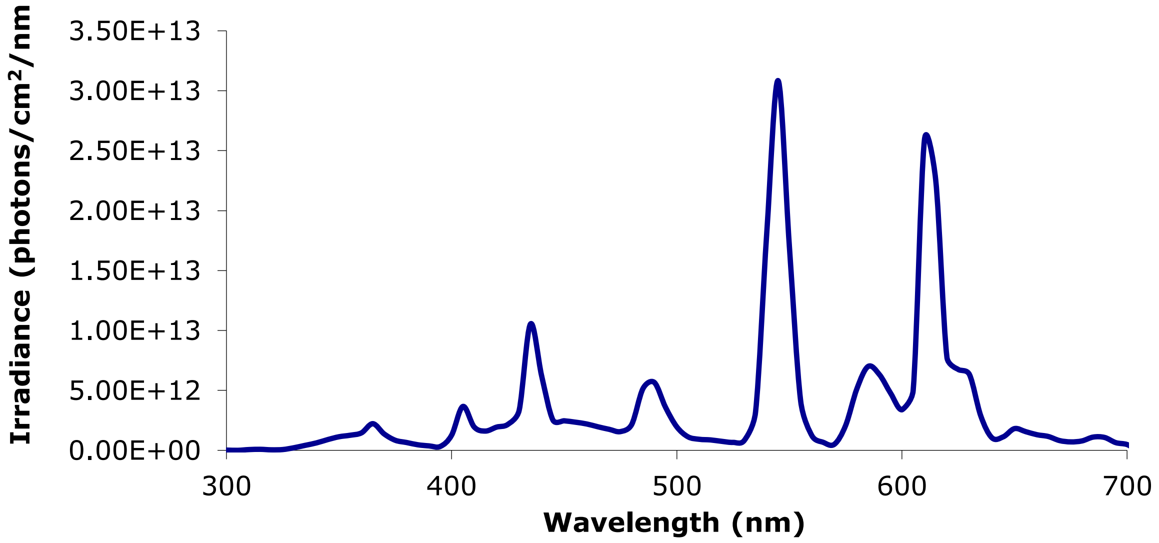

*Supplementary Table 1*. The location and read direction of mapped opsin genes in ten anemonefish species and a representative damselfish sister species (*Pomacentrus moluccensis*). For some species, it was not possible to map the full coding sequence, and only individual exons or partial exons were included.

| Species | Gene | Location (contig/chromosome no.) | Read direction (forward/reverse) |
|---|---|---|---|
| *A. akallopisos* | *SWS1β* | Contig 2050:<br>Exon 1: 5,580 - 5,244<br>Exon 2: 5,162 – 4,993<br>Exon 3: 2,667 – 2,499<br>Exon 4: 2,038 – 1,802<br>Exon 5: 810 – 706 | Reverse |
| | *SWS1α* | Contig 33, 668<br>Exon 3: 3,317 – 3,486<br>Exon 4: 4,185 – 4,422<br>Exon 5: 5,161 – 5,262 | Forward |
| | *SWS2B* | Contig 359: 279, 141 – 282, 538 | Reverse |
| | *RH2B* | Contig 29424: 10, 925- 13, 771 | Reverse |
| | *RH2A-2* | Contig 34,813:<br>Exon 1: 4,491 – 4,877<br>Part of Exon 2: 4,982 – 5,005 | Forward |
| | *RH2A-1* | Contig 34,450:<br>Exon 1: 113 – 499<br>Exon 2: 605 – 774<br>Exon 3: 961 – 1,124<br>Exon 4: 1,256 – 1,494<br>Exon 5: 1,600 – 1,696 | Forward |
| | *LWS* | Contig 359: 268,513- 271,073 | Reverse |

| | | | |
|---|---|---|---|
| | *RH1* | Contig 690: 246,742- 247,800 | Reverse |
| *A. bicinctus* | *SWS1α* | Contig 33,668<br>Exon 3: 2,153 – 2,322<br>Exon 4: 4,186 – 4,423<br>Exon 5: 5,164 – 5,265 | Forward |
| | *SWS1β* | Contig 2,050<br>Exon 1: 10, 184 – 10, 526<br>Exon 2: 10, 607 – 10, 776<br>Exon 3: 13, 101 – 13, 266<br>Exon 4: 13, 727 – 13, 964<br>Exon 5: 14, 956 – 15, 057 | Forward |
| | *SWS2B* | Contig 359: 278,952 - 282,349 | Reverse |
| | *RH2B* | Contig 29424: 10,869 - 13,715 | Reverse |
| | *RH2A-1* | Contig 34,450<br>Exon 1: 93 – 479<br>Exon 2: 585 – 754<br>Exon 3: 941 – 1,104<br>Exon 4: 1,236 – 1,474<br>Exon 5: 1,580 – 1,676 | Forward |
| | *RH2A-2* | Contig 34,813<br>Exon 1: 4,492 – 4,881 | Forward |
| | *LWS* | Contig 359: 268, 336 – 270, 884 | Reverse |
| | *RH1* | Contig 690: 246, 678 – 247, 736 | Reverse |
| *A. melanopus* | *SWS1β* | Contig 2, 050<br>Exon 1: 10, 212 – 10, 553<br>Exon 2: 10, 635 – 10, 804<br>Exon 3: 13, 128 – 13, 293 | Forward |

| | | | |
|---|---|---|---|
| | SWS1α | Exon 4: 13, 754 – 13, 991<br>Exon 5: 14, 983 – 15, 084 | |
| | | Contig 33, 668<br>Exon 4: 4, 206 – 4, 445<br>Exon 5: 5, 184 – 5, 285 | Forward |
| | *SWS2B* | Contig 359: 279, 206 – 282, 603 | Reverse |
| | *RH2B* | Contig 29424: 10, 874 - 13, 720 | Reverse |
| | *RH2A-1* | Contig 34, 450:<br>Exon 1: 309 – 695<br>Exon 2: 801- 970<br>Exon 3: 1, 157 – 1, 320<br>Exon 4: 1, 452 – 1, 690<br>Exon 5: 1, 796 – 1, 892 | Forward |
| | *RH2A-2* | Contig 34, 813<br>Exon 1: 4, 493 | Forward |
| | *LWS* | Contig 359: 268, 642 – 271, 138 | Reverse |
| | *RH1* | Contig 690: 247, 256 – 248, 314 | Reverse |
| *A. nigripes* | *SWS1β* | Contig 2, 050<br>Exon 1: 10, 195 – 10, 536<br>Exon 2: 10, 642 – 10, 787<br>Exon 3: 13, 111 – 13, 276<br>Exon 4: 13, 737 – 13, 974<br>Exon 5: 14, 966 – 15, 067 | Forward |
| | *SWS1α* | Contig 33, 668<br>Exon 4: 4, 205 – 4, 444<br>Exon 5: 5, 183 – 5, 284 | Forward |
| | *SWS2B* | Contig 359: 279, 709 – 283, 106 | Reverse |

| | | |
|---|---|---|
| *RH2B* | Contig 29424: 10, 923 – 13, 769 | Reverse |
| *RH2A-1* | Contig 34, 450<br>Exon 1: 103 – 489<br>Exon 2: 595 – 764<br>Exon 3: 951 – 1, 114<br>Exon 4: 1, 246 – 1, 484<br>Exon 5: 1, 590 – 1, 686 | Forward |
| *RH2A-2* | Contig 34, 813<br>Exon 1: 4, 491 – 4, 880 | Forward |
| *LWS* | Contig 359: 268, 834 – 271, 390 | Reverse |
| *RH1* | Contig 690: 246, 674 – 247, 732 | Reverse |

| | | | |
|---|---|---|---|
| *A. Perideraion* | *SWS1β* | Contig 2,050<br>Exon 1: 10, 549 – 10, 890<br>Exon 2: 10, 972 – 11, 141<br>Exon 3: 13, 466 – 13, 631<br>Exon 4: 14, 092 – 14, 329<br>Exon 5: 15, 322 - 15, 423 | Forward |
| | *SWS1α* | Contig 33, 668<br>Exon 3: 3, 515 – 3, 684<br>Exon 4: 4, 149 – 4, 386<br>Exon 5: 5, 125 – 5, 226 | Forward |
| | *SWS2B* | Contig 359: 279, 488- 282, 885 | Reverse |
| | *RH2B* | Contig 29424: 10, 919 – 13, 765 | Reverse |
| | *RH2A-1* | Contig 34450<br>Exon 1: 91- 477 | Forward |

| | | | |
|---|---|---|---|
| | | Exon 2: 583 – 752<br>Exon 3: 939 – 1, 102<br>Exon 4: 1, 234 – 1, 472<br>Exon 5: 1, 578 – 1, 674 | |
| | *RH2A-2* | Contig 34, 813<br>Exon 1: 4, 492 – 4, 878<br>Part of Exon 2: 4, 983 – 5,004 | Forward |
| | *LWS* | Contig 359: 268, 685 – 271, 177 | Reverse |
| | *RH1* | Contig 690: 246, 842 – 247, 900 | Reverse |
| *A. Polymnus* | *SWS1β* | Contig 2050:<br>Exon 1: 9, 998 – 10, 339<br>Exon 2: 10, 430 – 10, 590<br>Exon 3: 12, 914 – 13, 079<br>Exon 4: 13, 540 – 13, 777<br>Exon 5: 14, 769 – 14, 870 | Forward |
| | *SWS1α* | Contig 33, 668<br>Exon 4: 4, 207 – 4, 446<br>Exon 5: 5, 186 – 5, 287 | Forward |
| | *SWS2B* | Contig 359: 279, 123 – 282, 520 | Reverse |
| | *RH2B* | Contig 29424: 10, 861 – 13, 707 | Reverse |
| | *RH2A-1* | Contig 34, 450<br>Exon 1: 112 – 498<br>Exon 2: 604 – 773<br>Exon 3: 960 – 1, 123<br>Exon 4: 1, 255 – 1, 493<br>Exon 5: 1, 599 – 1, 695 | Forward |
| | *RH2A-2* | Contig 34, 813 | Forward |

| | | | |
|---|---|---|---|
| | | Exon 1: 4, 491 – 4, 880 | |
| | *LWS* | Contig 359: 268, 262 – 270, 806 | Reverse |
| | *RH1* | Contig 690: 246, 975- 248, 033 | Reverse |
| *A. sebae* | *SWS1β* | Contig 2050<br>Exon 1: 9,992 – 10, 333<br>Exon 2: 10, 424 – 10, 584<br>Exon 3: 12, 910 – 13, 075<br>Exon 4: 13, 536 – 13, 773<br>Exon 5: 14, 765 – 14, 866 | Forward |
| | *SWS1α* | Contig 33, 668<br>Exon 3: 1, 585 – 1, 824<br>Exon 4: 2, 570 – 2, 625<br>Exon 5: 5, 240 – 5, 285 | Forward |
| | *SWS2B* | Contig 359: 278, 944 – 282, 341 | Reverse |
|  | *RH2B* | Contig 29424: 10, 867 – 13, 713 | Reverse |
| | *RH2A-1* | Contig 34, 450<br>Exon 1: 103 – 489<br>Exon 2: 595 – 764<br>Exon 3: 951 – 1, 114<br>Exon 4: 1, 246 – 1, 484<br>Exon 5: 1, 590 – 1, 686 | Forward |
| | *RH2A-2* | Contig 34, 813:<br>Exon 1: 4, 491 – 4, 880 | Forward |
| | *LWS* | Contig 359: 268, 082 – 270, 626 | Reverse |
| | *RH1* | Contig 690: 246, 855 – 247, 913 | Reverse |
| *P. Moluccensis* | | Contig 2050: | Forward |

| | | | |
|---|---|---|---|
| | *SWS1β* | Exon 1: 10, 198 – 10, 539<br>Exon 2: 10, 634 - 10, 790<br>Exon 3: 13, 114 – 13, 279<br>Exon 4: 13, 740 – 13, 983<br>Exon 5: 14, 976 – 15 | |
| | *SWS1α* | Contig 33, 668<br>Exon 4: 1, 814 – 2, 059<br>Exon 5: 5, 399 – 5, 494 | Forward |
| | *SWS2B* | Contig 359: 278, 007 – 281, 404 | Reverse |
| | *RH2B* | Contig 29424: 10, 849 – 13, 695 | Reverse |
| | *RH2A-1* | Contig 34, 450:<br>Exon 1: 316 – 702<br>Exon 2: 808 – 977<br>Exon 3: 1, 164 – 1, 327<br>Exon 4: 1, 459 – 1, 697<br>Exon 5: 1, 803 – 1, 899 | Forward |
| | *RH2A-2* | Contig 34, 494:<br>Exon 1: 4, 494 – 4, 880<br>Part of Exon 2: 4, 985 – 5, 023 | Forward |
| | *LWS* | Conti 359: 267, 421 – 269, 938 | Reverse |
| | *RH1* | Contig 690: 246, 745 – 247, 803 | Reverse |
| *P. biaculeatus* | *SWS1β* | Contig 2050:<br>Exon 1: 5243 – 5582<br>Exon 2: 4993 – 5161<br>Exon 3: 2501 – 2667<br>Exon 4: 1801 – 2040 | Reverse |

| | | | |
|---|---|---|---|
| | | Exon 5: 706 – 807 | |
| | SWS1α | Contig 33668:<br>Exon 4: 1801 – 2040<br>Exon 5: 706 – 807 | Reverse |
| | SWS2B | Contig 359: 279, 037 – 282, 434 | Reverse |
| | RH2B | Contig 29424: 10, 844 – 13, 690 | Reverse |
| | RH2A-1 | Contig 34, 450:<br>Exon 1: 604 – 988<br>Exon 2: 1, 094 – 1, 262<br>Exon 3: 1, 449 – 1, 614<br>Exon 4: 1, 746 – 1, 985<br>Exon 5: 2, 091 – 2, 187 | Reverse |
| | RH2A-2 | Contig 34, 813:<br>Exon 1: 169 – 553 | Reverse |
| | LWS | Contig 359: 268, 225 – 270, 721 | Reverse |
| | RH1 | Contig 690: 242, 934 – 243, 992 | Reverse |
| *A. frenatus* | SWS1β | Contig 2050:<br>Exon 1: 5, 239 - 5, 579<br>Exon 2: 4, 989 – 5, 157<br>Exon 3: 2, 499 – 2, 665<br>Exon 4: 1, 799 – 2, 038<br>Exon 5: 706 – 807 | Reverse |
| | SWS1α | Contig 33, 668:<br>Exon 4: 21, 239 – 21, 478 | Reverse |

| | | | |
|---|---|---|---|
| | | Exon 5: 17, 801 – 17, 902 | |
| | *SWS2B* | Contig 359: 278, 535 – 281, 932 | Reverse |
| | *RH2B* | Contig 29424: 10, 785 – 13, 631 | Reverse |
| | *RH2A-1* | Contig 34, 450:<br>Exon 1: 671 – 1, 055<br>Exon 2: 1, 161 – 1, 329<br>Exon 3: 1, 516 – 1, 681<br>Exon 4: 1, 813 – 2, 052<br>Exon 5: 2, 158 – 2, 256 | Forward |
| | *RH2A-2* | Contig 34, 813:<br>Exon 1: 199 - 583 | Reverse |
| | *LWS* | Contig 359: 267, 952 – 269, 300 | Reverse |
| | *RH1* | Contig 690: 246, 539 – 247, 597 | Reverse |
| *A. ocellaris* | *SWS1β* | Contig 2, 110:<br>Exon 1: 7, 017 – 7, 438<br>Exon 2: 6, 849 – 7. 017<br>Exon 3: 4, 408 – 4, 573<br>Exon 4: 3, 704 – 3, 943<br>Exon 5: 2, 592 – 2, 693 | Reverse |
| | *SWS1α* | Contig 4, 729:<br>Exon 1: 8, 709 – 9, 048<br>Exon 2: 9, 130 – 9, 208<br>Exon 3: 10, 474 – 10, 639<br>Exon 4: 12, 839 – 13, 078 | Forward |

| | | | |
|---|---|---|---|
| | | Exon 5: 13, 887 – 13, 988 | |
| | *SWS2B* | Contig 2946: 168, 673 – 171, 978 | Forward |
| | *RH2B* | Contig 3040: 608, 700 – 611. 484 | Reverse |
| | *RH2A-1* | Contig 3040:<br>Exon 1: 583, 563 – 583, 947<br>Exon 2: 583, 289 – 583, 457<br>Exon 3: 582, 942 – 583, 107<br>Exon 4: 582, 576 – 582, 815<br>Exon 5: 582, 372 – 582, 470 | Reverse |
| | *RH2A-2* | Contig 3040:<br>Exon 1: 593, 798 – 594, 182<br>Exon 2: 593, 524 – 593, 692<br>Exon 3: 593, 144 – 593, 309<br>Exon 4: 592, 778 – 593, 017<br>Exon 5: 592, 574 – 592, 672 | Reverse |
| | *LWS* | Contig 2946: 179, 072 - 182, 064 | Forward |
| | *RH1* | Contig 3183: 242, 934 – 243, 992 | Forward |
| *A. percula* | *SWS1β* | Chromosome 21:<br>Exon 1: 29, 064, 435 – 29, 064, 774<br>Exon 2: 29, 064, 185 – 29, 064, 353<br>Exon 3: 29, 061, 743 – 29, 061, 908<br>Exon 4: 29, 061, 043 – 29, 061, 282<br>Exon 5: 29, 059, 944 – 29, 060, 045 | Reverse |
| | | Chromosome 21:<br>Exon 1: 29, 040, 288 – 29, 040, 627 | Reverse |

| | | |
|---|---|---|
| *SWS1α* | Exon 2: 29, 040, 038 – 29, 040, 206<br><br>Exon 3: 29, 038, 703 – 29, 038, 868<br><br>Exon 4: 29, 036, 681 – 29, 036, 920<br><br>Exon 5: 29, 035, 775 – 29, 035, 876 | |
| *SWS2B* | Chromosome 6: 20, 661, 387 – 20, 664, 781 | Reverse |
| *RH2B* | Chromosome 6: 11, 995, 019 – 11, 997, 839 | Reverse |
| *RH2A-1* | Chromosome 6: 11, 968, 126 – 11, 969, 712 | Reverse |
| *RH2A-2* | Chromosome 6: 11, 978, 627 – 11, 980, 238 | Reverse |
| *RH1* | Chromosome 8: 12, 259, 415 – 12, 260, 473 | Reverse |

*Supplementary Table 2*. The location and read direction of visual transduction pathway genes in *A. percula* and *A. ocellaris*. Predicted gene sequences were obtained from reference annotations openly available on Ensembl (ensembl.org). Coding sequences were verified by assembly against the transcriptome of the *A. ocellaris* retina.

| Species | Gene | Location (contig/chromosome no.) | Read direction (forward/reverse) |
|---|---|---|---|
| *A. percula* | *cnga3* | Chromosome 2: 9,749,530-9,757,646 | Reverse |
| | *cngb3-1* | Chromosome 22: 25,237,165-25,251,431 | Reverse |
| | *cngb3-2* | Chromosome 22: 25,254,499-25,261,458 | Reverse |
| | *gnat1* | Chromosome 6: 5,339,816-5,343,098 | Forward |
| | *gnat2* | Chromosome 8: 5,554,610-5,562,717 | Reverse |
| | *gnb1* | Chromosome 8: 27,879,688-27,914,153 | Reverse |
| | *gnb2* | Chromosome 15: 18,211,100-18,223,225 | Forward |
| | *gnb3* | Chromosome 7: 36,912,225-36,921,246 | Forward |
| | *pde6c* | Chromosome 19: 30,171,117-30,185,145 | Forward |
| | *pde6h* | Chromosome 3: 29,046,539-29,048,777 | Reverse |
| *A. ocellaris* | *cnga3* | NXFZ01003666.1: 88,031-95,226 | Forward |
| | *cngb3-1* | NXFZ01005675.1: 602,781-619,472 | Forward |
| | *cngb3-2* | NXFZ01005675.1: 593,521-600,562 | Forward |
| | *gnat1* | NXFZ01004316.1: 227,748-230,552 | Forward |
| | *gnat2* | NXFZ01003723.1: 3,374-8,842 | Forward |
| | *gnb1* | NXFZ01002376.1: 1,540,561-1,560,162 | Reverse |
| | *gnb2* | NXFZ01004072.1: 39,382-44,533 | Forward |
| | *gnb3* | NXFZ01002559.1: 34,228-38,944 | Forward |
| | *pde6c* | NXFZ01002893.1: 580,901-581,368 | Forward |
| | *pde6h* | NXFZ01003767.1: 210,269-210,608 | Forward |

*Supplementary Table 3*. The location and read direction of visual transduction shutoff genes in *A. percula* and *A. ocellaris*. Predicted gene sequences were obtained from annotations openly available on Ensembl (ensembl.org). All coding sequences were verified by assembly against the transcriptome of the *A. percula* and *A. ocellaris* retina.

| Species | Gene | Location (contig/chromosome no.) | Read direction (forward/reverse) |
|---|---|---|---|
| | *arr3* | Chromosome 13: 22,323,604-28,328,645 | Forward |
| | *arrb1* | Chromosome 9: 38,538,861-38,548,645 | Reverse |
| | *arrb2* | Chromosome 15: 1,003,671-1,029,284 | Reverse |
| | *gnb5* | Chromosome 4: 15,152,646-15,157,681 | Reverse |
| *A. percula* | *grk1* | Chromosome 12: 37,918,601-37,924,758 | Reverse |
| | *grk7* | Chromosome 10: 24,358,798-24,366,631 | Reverse |
| | *rgs11* | Chromosome 10: 29,326,188-29,336,761 | Forward |
| | *rgs6* | Chromosome 11: 19,343,058-19,373,376 | Forward |
| | *rgs7* | Chromosome 20: 6,476,716-6,529,039 | Reverse |
| | *rgs9* | Chromosome 14: 84,740-108,809 | Reverse |
| | *rgs9bp* | Chromosome 22: 24,147,177-24,152,121 | Forward |
| | *sag* | Chromosome 9: 35,513,569-35,525,733 | Reverse |
| | *arr3* | NXFZ1002075.1: 95,106-97,935 | Reverse |
| | *arrb1* | NXFZ1006210.1: 155,747-167,318 | Reverse |
| | *arrb2* | NXFZ01004797.1: 12-28,906 | Forward |
| | *gnb5* | NXFZ01003222.1: 87,542-91,935 | Reverse |
| | *grk1* | NXFZ01000019.1: 428,677-438,372 | Reverse |
| *A. ocellaris* | *grk7* | NXFZ01004526.1: 124,442-130,432 | Reverse |
| | *rgs11* | NXFZ01001898.1: 1,268,205-1,278,181 | Reverse |
| | *rgs6* | NXFZ01002380.1: 170,082-200,306 | Forward |
| | *rgs7* | NXFZ01003727.1: 491,552-515,327 | Reverse |
| | *rgs9* | NXFZ01002076.1: 61,111-76,301 | Forward |
| | *rgs9bp* | NXFZ01001919.1: 840,381-844,179 | Forward |
| | *sag* | NXFZ01004064.1: 81,804-95,808 | Reverse |

Supplementary Figure 2. Variable amino acid sites and estimated tuning effects in anemonefish cone opsins including LWS, RH2A, RH2B, SWS2B and SWS1. Opsin protein alignment and amino acid site number corresponds to a bovine rhodopsin template (PDB accession no. 1U19). Estimated tuning effects on opsin λmax of amino acid substitutions were either based on in-vitro measurements (see cited sources) or deduced from differences between the reference sequences and their λmax values. Opsin λmax value estimates are listed in the final column with the reference sequence that calculations were based in parentheses. An exact match in known tuning sites were found for both SWS1 opsins in A. percula and A. ocellaris (shown in bold).

| Species | Variable sites | | | Estimated tuning effec | | | | Actual |
|---|---|---|---|---|---|---|---|---|
| Anemonefishes | A40S | S164A | Y178F | A40S | S164A | Y178F | Estimate (nm) | $\lambda_{max}$ (nm) |
| LWS | | | | | | | | |
| Oreochromis niloticus | A | S | F | | | | 561 | 561 |
| Oryzias latipes | A | S | Y | 0 | 0 | -1 | 560 | 560 |
| Lucania goodei a | S | S | Y | 13 | 0 | -1 | 573 | 573 |
| Maylandia zebra | A | A | Y | 0 | -7 | -1 | 554 | 554 |
| Amphiprion percula | A | S | Y | 0 | 0 | -1 | 560 (O. niloticus, O. latipes, M. zebra ), 561 ( L. goodei) | |
| Amphiprion ocellaris | A | S | Y | 0 | 0 | -1 | 560 (O. niloticus, O. latipes, M. zebra ), 561 ( L. goodei) | |
| Amphiprion frenatus | A | S | Y | 0 | 0 | -1 | 560 (O. niloticus, O. latipes, M. zebra ), 561 ( L. goodei) | |
| Premnas biaculeatus | A | S | Y | 0 | 0 | -1 | 560 (O. niloticus, O. latipes, M. zebra ), 561 ( L. goodei) | |
| Amphiprion nigripes | A | A | Y | 0 | -7 | -1 | 554 (M. zebra ), 553 ( O. niloticus, L. goodei ), 561 ( O. latipes ) | |
| Amphiprion polymnus | A | A | Y | 0 | -7 | -1 | 554 (M. zebra ), 553 ( O. niloticus, L. goodei ), 561 ( O. latipes ) | |
| Amphiprion perideraion | A | S | Y | 0 | 0 | -1 | 560 (O. niloticus, O. latipes, M. zebra ), 561 ( L. goodei) | |
| Amphiprion akallopisos | A | S | Y | 0 | 0 | -1 | 560 (O. niloticus, O. latipes, M. zebra ), 561 ( L. goodei) | |
| Amphiprion sebae | A | A | Y | 0 | -7 | -1 | 554 (M. zebra ), 553 ( O. niloticus, L. goodei ), 561 ( O. latipes ) | |
| Amphiprion bicinctus | A | A | Y | 0 | -7 | -1 | 554 (M. zebra ), 553 ( O. niloticus, L. goodei ), 561 ( O. latipes ) | |
| Amphiprion melanopus | A | S | Y | 0 | 0 | -1 | 560 (O. niloticus, O. latipes, M. zebra ), 561 ( L. goodei) | |
| | | S164A = -7<br>A164S = 6<br>Yokoyama, 2008 | | | | | | |

| Species | | | | | | | | | Estimated tuning effect | | | |
|---|---|---|---|---|---|---|---|---|---|---|---|---|
| Anemonefishes | P107S/T | F109N/A | S124A | Y37F | F60L/Y | V255I | T266V/K | F158L/I | P107S/T | F158L/I/T | Estimate (nm) | Actual $\lambda_{max}$ (nm) |
| RH2A | | | | | | | | | | | | |
| Oreochromis niloticus RH2A-1 | P | F | A | Y | F | V | T | F | | | 528 | 528 |
| Oreochromis niloticus RH2A-2 | T | F | A | F | F | V | T | L | 0 | -10 | 518 | 518 |
| Pomacentrus amboinensis | P | F | A | Y | F | V | T | I | 0 | -10 | 518 | 523 |
| Maylandia zebra RH2A-1 | P | F | A | Y | F | V | T | F | 0 | 0 | 528 | 528 |
| Maylandia zebra RH2A-2 | S | F | A | F | F | V | T | L | 1 | -10 | 518 | 519 |
| Dascyllus trimaculatus | P | F | A | Y | F | V | T | I | 0 | -10 | 518 | 516 |
| Amphiprion percula RH2A-1 | P | F | A | Y | F | V | T | I | 0 | -10 | 518 (O. niloticus RH2A-1 , M. zebra RH2A-1 , 523 (P. amboinensis, 516 (D. trimaculatus ) | |
| Amphiprion percula RH2A-2 | P | A | S | F | F | V | V | I | 0 | -10 | 518 (O. niloticus RH2A-1 , M. zebra RH2A-1 ), 523 ( P. amboinensis), 516 ( D. trimaculatus ) | |
| Amphiprion ocellaris RH2A-1 | P | F | A | Y | F | V | T | I | 0 | -10 | 518 (O. niloticus RH2A-1 , M. zebra RH2A-1 , 523 (P. amboinensis, 516 (D. trimaculatus ) | |
| Amphiprion ocellaris RH2A-2 | P | A | A | F | F | V | V | I | 0 | -10 | 518 (O. niloticus RH2A-1 , M. zebra RH2A-1 P. amboinensis D. trimaculatus ) | |
| Amphiprion frenatus RH2A-1 | P | F | A | Y | F | V | T | I | 0 | -10 | 518 (O. niloticus RH2A-1 , M. zebra RH2A-1 , 523 (P. amboinensis, 516 (D. trimaculatus ) | |
| Premnas biaculeatus RH2A-1 | P | F | A | Y | F | V | T | I | 0 | -10 | 518 (O. niloticus RH2A-1 , M. zebra RH2A-1, 523 ( P. amboinensis, 516 (D. trimaculatus ) | |
| Amphiprion nigripes RH2A-1 | P | F | A | Y | F | V | V | M | 0 | 0 | 528 (O. niloticus RH2A-1, M. zebra RH2A-1 ) | |
| Amphiprion polymnus RH2A-1 | P | F | A | Y | F | V | V | M | 0 | 0 | 528 (O. niloticus RH2A-1, M. zebra RH2A-1 ) | |
| Amphiprion perideraion RH2A-1 | P | F | A | Y | F | I | K | I | 0 | -10 | 518 (O. niloticus RH2A-1 , M. zebra RH2A-1 ), 523 ( P. amboinensis), 516 ( D. trimaculatus ) | |
| Amphiprion akallopisos RH2A-1 | P | N | A | Y | Y | I | K | I | 0 | -10 | 518 (O. niloticus RH2A-1 , M. zebra RH2A-1 ), 523 ( P. amboinensis), 516 ( D. trimaculatus ) | |
| Amphiprion sebae RH2A-1 | P | F | A | Y | F | V | V | M | 0 | 0 | 528 (O. niloticus RH2A-1, M. zebra RH2A-1 ) | |
| Amphiprion bicinctus RH2A-1 | P | F | A | Y | F | V | V | T | 0 | -10 | 518 (O. niloticus RH2A-1 , M. zebra RH2A-1 ), 523 ( P. amboinensis), 516 ( D. trimaculatus ) | |
| Amphiprion melanopus RH2A-1 | P | F | A | Y | F | V | T | I | 0 | -10 | 518 (O. niloticus RH2A-1 , M. zebra RH2A-1, 523 ( P. amboinensis, 516 (D. trimaculatus ) | |
| | | | | | | | | | | F158L/I = -10<br>Spady et al. 2006<br>Parry et al. 2005 | | |

| Species | | | | | | | | Estimated tuning effect | | | | | Actual |
|---|---|---|---|---|---|---|---|---|---|---|---|---|---|
| Anemonefishes | I49C | M44I | S109G | Q122K | M207L | A166S/A166T | S124A | M44I | S109G | Q122K | M207L | A166S/A166T | Estimated λmax (nm) | $\lambda_{max}$ (nm) |
| RH2B | | | | | | | | | | | | | | |
| Oreochromis niloticus | I | M | S | K | M | S | S | 0 | | 0 | 0 | 0 | | 472 |
| Pomacentrus amboinensis | C | M | G | K | M | S | S | 0 | 8 | 0 | 0 | 0 | 480 | 480 |
| Maylandia zebra | I | I | S | Q | M | A | S | 3 | 0 | 17 | 0 | -8 | 484 | 484 |
| Dascyllus trimaculatus | C | M | G | Q | L | T | S | 0 | 8 | 17 | -7 | 0 | 490 | 490 |
| Amphiprion percula | C | M | G | Q | M | T | A | 0 | 8 | 17 | 0 | 0 | 497 (O. niloticus, P. amboinensis, M. zebra, D. trimaculatus ) | |
| Amphiprion ocellaris | C | M | G | Q | M | T | A | 0 | 8 | 17 | 0 | 0 | 497 (O. niloticus, P. amboinensis, M. zebra, D. trimaculatus ) | |
| Amphiprion frenatus | C | M | G | Q | M | T | A | 0 | 8 | 17 | 0 | 0 | 497 (O. niloticus, P. amboinensis, M. zebra, D. trimaculatus ) | |
| Premnas biaculeatus | C | M | G | Q | M | T | A | 0 | 8 | 17 | 0 | 0 | 497 (O. niloticus, P. amboinensis, M. zebra, D. trimaculatus ) | |
| Amphiprion nigripes | C | M | G | Q | M | T | S | 0 | 8 | 17 | 0 | 0 | 497 (O. niloticus, P. amboinensis, M. zebra, D. trimaculatus ) | |
| Amphiprion polymnus | C | M | G | Q | M | T | S | 0 | 8 | 17 | 0 | 0 | 497 (O. niloticus, P. amboinensis, M. zebra, D. trimaculatus ) | |
| Amphiprion perideraion | C | M | G | Q | M | T | S | 0 | 8 | 17 | 0 | 0 | 497 (O. niloticus, P. amboinensis, M. zebra, D. trimaculatus ) | |
| Amphiprion akallopisos | C | M | G | Q | M | T | A | 0 | 8 | 17 | 0 | 0 | 497 (O. niloticus, P. amboinensis, M. zebra, D. trimaculatus ) | |
| Amphiprion sebae | C | M | G | Q | M | T | S | 0 | 8 | 17 | 0 | 0 | 497 (O. niloticus, P. amboinensis, M. zebra, D. trimaculatus ) | |
| Amphiprion bicinctus | C | M | G | Q | M | T | S | 0 | 8 | 17 | 0 | 0 | 497 (O. niloticus, P. amboinensis, M. zebra, D. trimaculatus ) | |
| Amphiprion melanopus | C | M | G | Q | M | T | S | 0 | 8 | 17 | 0 | 0 | 497 (O. niloticus, P. amboinensis, M. zebra, D. trimaculatus ) | |
| | | | | | | | | M44I = +3<br>Yokoyama & Jia, 2020 | S109G = +8 when I49C present<br>Luehrmann et al. 2019 | Q122K (+15)<br>Q122E (+17)<br>Yokoyama & Jia, 2020 | M207L = -7<br>Yokoyama & Jia, 2020 | Suspected tuning site<br>Yokoyama & Jia, 2020 | | |

| Species | Variable sites | | | | | | Estimated tuning effect | | | | | | | Actual |
|---|---|---|---|---|---|---|---|---|---|---|---|---|---|---|
| Anemonefishes | V46F | A109G | T118A/G | A168S/T | Y203F | Y265W | V46F | A109G | S/T118A/G | A168S/T | Y203F | Y265W | Estimate (nm) | $\lambda_{max}$ (nm) |
| SWS2B | | | | | | | | | | | | | | |
| Oreochromis niloticus | F | A | T | A | Y | W | | | | | | | 423 | 423 |
| Maylandia zebra | F | A | T | A | Y | W | 0 | 0 | 0 | 0 | 0 | 0 | 423 | 423 |
| Oryzias latipes | V | G | A | S | F | Y | 6 | -2 | -15 | 6 | 1 | -15 | 404 | 405 |
| Lucania goodei | F | G | G | T | Y | Y | 0 | -2 | -15 | 6 | 0 | -15 | 397 | 397 |
| Amphiprion percula | F | G | T | A | F | Y | 0 | -2 | 0 | 0 | 1 | -15 | 407 ( O . niloticus , M. zebra, O. latipes, L. goodei ) | |
| Amphiprion ocellaris | F | G | T | A | F | Y | 0 | -2 | 0 | 0 | 1 | -15 | 407 ( O . niloticus , M. zebra, O. latipes, L. goodei ) | |
| Amphiprion frenatus | F | G | T | A | Y | Y | 0 | -2 | 0 | 0 | 0 | -15 | 406 ( O . niloticus , M. zebra, O. latipes, L. goodei ) | |
| Premnas biaculeatus | F | G | T | A | F | Y | 0 | -2 | 0 | 0 | 1 | -15 | 407 ( O . niloticus , M. zebra, O. latipes, L. goodei ) | |
| Amphiprion nigripes | F | G | S | A | Y | Y | 0 | -2 | 0 | 0 | 0 | -15 | 406 ( O . niloticus , M. zebra, O. latipes, L. goodei ) | |
| Amphiprion polymnus | F | G | S | A | Y | Y | 0 | -2 | 0 | 0 | 0 | -15 | 406 ( O . niloticus , M. zebra, O. latipes, L. goodei ) | |
| Amphiprion perideraion | F | G | S | A | Y | Y | 0 | -2 | 0 | 0 | 0 | -15 | 406 ( O . niloticus , M. zebra, O. latipes, L. goodei ) | |
| Amphiprion akallopisos | F | G | S | A | Y | Y | 0 | -2 | 0 | 0 | 0 | -15 | 406 ( O . niloticus , M. zebra, O. latipes, L. goodei ) | |
| Amphiprion sebae | F | G | S | A | Y | Y | 0 | -2 | 0 | 0 | 0 | -15 | 406 ( O . niloticus , M. zebra, O. latipes, L. goodei ) | |
| Amphiprion bicinctus | F | G | S | A | Y | Y | 0 | -2 | 0 | 0 | 0 | -15 | 406 ( O . niloticus , M. zebra, O. latipes, L. goodei ) | |
| Amphiprion melanopus | F | G | S | A | Y | Y | 0 | -2 | 0 | 0 | 0 | -15 | 406 ( O . niloticus , M. zebra, O. latipes, L. goodei ) | |
| | | | | | | | L46F (SWS2) = -6<br>Yokoyama et al 2007 | A109G (SWS2) = -2<br>Yokoyama et al 2007 | T118G (SWS2) = -15<br>Yokoyama et al. 2007<br>T118A (RH1) = -16<br>Janz & Farrens, 2001 | A164S (M/LWS) = +6<br>Neitz et al. 1991 | Y203F (SWS2) = +1<br>Estimated effect by Carleton et al. 2005 | W265Y (SWS2) = -29<br>Yokoyama et al. 2007<br>Y265W (SWS1) = +10<br>Fasick et al. 1999<br>W265Y (RH1) = -15<br>Nakamaya & Khorana, 1991<br>Nagata et al. 2002 | | |

| Species | Variable sites | | | | | | Estimated tuning effect | | | Actual |
|---|---|---|---|---|---|---|---|---|---|---|
| Anemonefishes | F49C | S82A | A118S | A125S | S168A | A114S | A118S | A114S | Estimate (nm) | $\lambda_{max}$ (nm) |
| SWS1 | | | | | | | | | | |
| Oreochromis niloticus | F | S | A | A | A | A | 0 | 0 | | 360 |
| Maylandia zebra | F | S | S | A | A | S | 5 | 3 | 368 | 368 |
| Lucania goodei | F | A | A | S | S | A | 0 | 0 | 360 | 354 |
| Pomacentrus amboinensis | C | A | S | A | A | S | 5 | 3 | 368 | 370 |
| Oryzias latipes | F | A | A | S | A | A | 0 | 0 | 360 | 356 |
| Amphiprion ocellaris SWS1a | F | A | A | S | A | A | 0 | 0 | 360 (O. niloticus , M. zebra ), 362 ( P. amboinensis), 356 (O. latipes | |
| Amphiprion ocellaris SWS1b | C | A | S | A | A | S | 5 | 3 | 363 (O. niloticus , M. zebra ), 370 (P. amboinensi) | |
| Amphiprion percula SWS1a | F | A | A | S | A | A | 0 | 0 | 360 (O. niloticus , M. zebra ), 362 ( P. amboinensis), 356 (O. latipes | |
| Amphiprion percula SWS1b | C | A | S | A | A | S | 5 | 3 | 363 (O. niloticus , M. zebra ), 370 (P. amboinensi) | |
| Amphiprion frenatus SWS1b | F | A | A | S | A | A | 0 | 0 | 360 (O. niloticus , M. zebra ), 362 ( P. amboinensis), 356 (O. latipes | |
| | | | | | | | A118S = +5<br>Shi & Yokoyama, 2003 | | | |

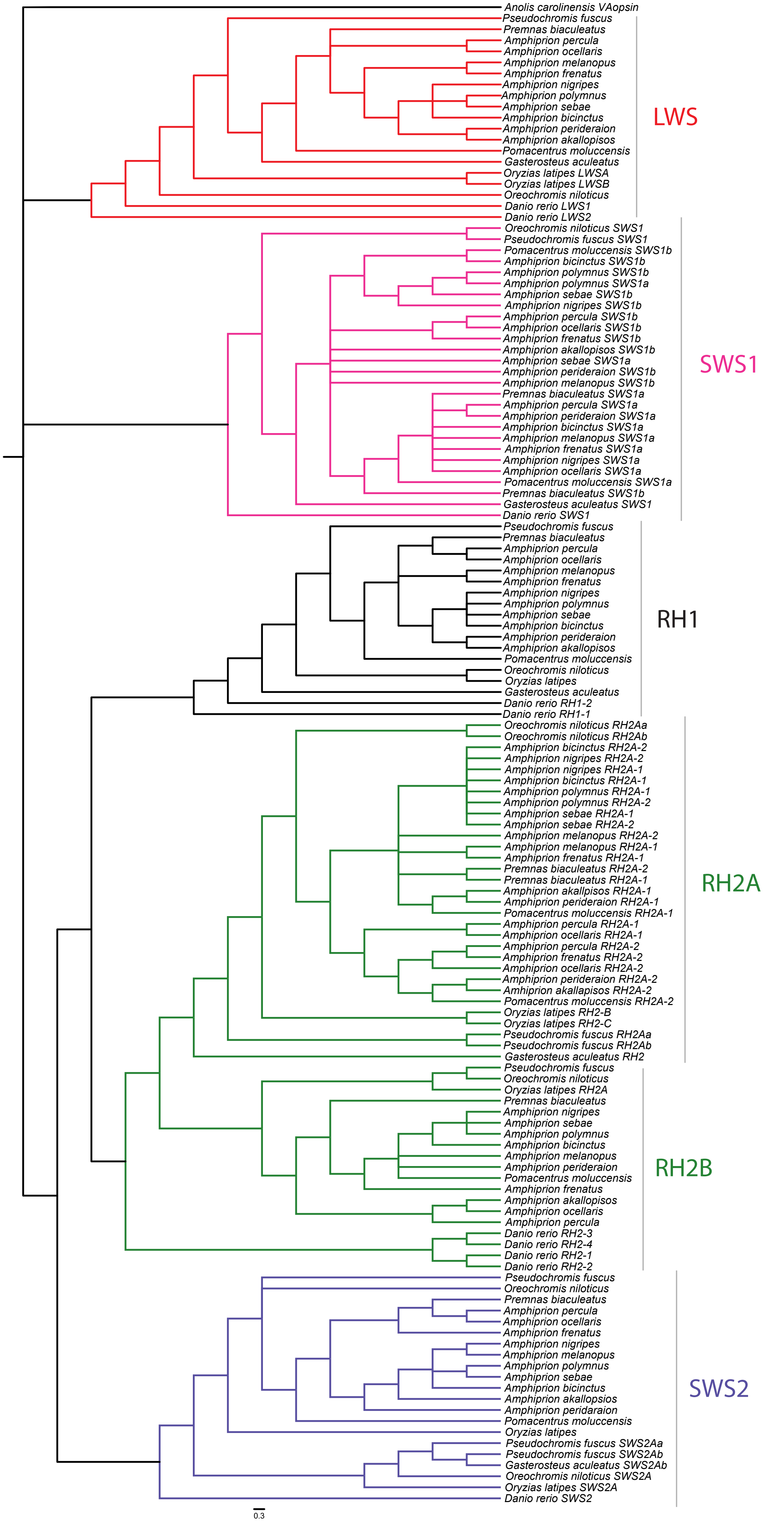

Supplementary Figure 3. Anemonefish visual opsin gene phylogeny reconstructed using the full-coding regions of all opsin genes. Shown are internodal Bayesian posterior probabilities (P) depicted as 'dark grey' and 'light grey' markers that indicate posterior probabilities greater than 0.95 and 0.9, respectively. Posterior probabilities less than 0.9 are given in full. Opsin gene acronyms stand for RH1 = Rhodopsin 1 (rod opsin), RH2 = Rhodopsin-like 2, SWS2 = Short-wavelength-sensitive 2, LWS = Long-wavelength-sensitive, SWS1 = Short-wavelength-sensitive 1, va = vertebrate ancient opsin (outgroup).

*Supplementary Figure 4*. Modelled light absorption curve for coexpression of *RH2A-1* (40%) with *LWS* (60%) ($\lambda_{max}$ value = 537-539 nm). Plotted alongside for comparison are modelled spectral absorbance curves for lone visual pigments and actual measured cone absorbance spectra.

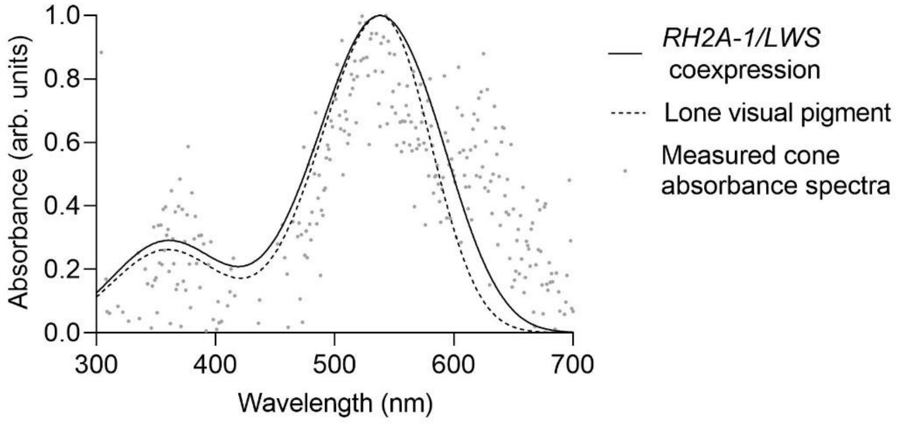

**Jmodel test output**

\-------------------------------------------------------------

*                                              *

*           AKAIKE INFORMATION CRITERION (AIC)           *

*                                              *

\-------------------------------------------------------------

Model selected:

Model = GTR+I+G

partition = 012345

-lnL = 48337.6631

K = 268

freqA = 0.2663

freqC = 0.2220

freqG = 0.2014

freqT = 0.3103

R(a) [AC] = 0.9051

R(b) [AG] = 1.9916

R(c) [AT] = 0.8833

R(d) [CG] = 1.0357

R(e) [CT] = 2.4097

R(f) [GT] = 1.0000

p-inv = 0.0460

gamma shape = 1.2620

\--

PAUP* Commands Block:

If you want to load the selected model and associated estimates in PAUP*,

attach the next block of commands after the data in your PAUP file:

[!

Likelihood settings from best-fit model (GTR+I+G) selected by AIC with jModeltest 2.1.10 v20160303 on Mon Jun 01 23:57:52 PDT 2020]

BEGIN PAUP;

Lset base=(0.2663 0.2220 0.2014 ) nst=6  rmat=(0.9051 1.9916 0.8833 1.0357 2.4097) rates=gamma shape=1.2620 ncat=4 pinvar=0.0460;

END;

--

* AIC MODEL SELECTION : Selection uncertainty

| Model | -lnL | K | AIC | delta | weight | cumWeight |
|---|---|---|---|---|---|---|
| GTR+I+G | 48337.66307 | 268 | 97211.326140 | 0.000000 | 0.999533 | 0.999533 |
| GTR+G | 48346.91648 | 267 | 97227.832960 | 16.506820 | 0.000260 | 0.999793 |
| HKY+I+G | 48350.14742 | 264 | 97228.294840 | 16.968700 | 0.000207 | 1.000000 |
| HKY+G | 48362.86273 | 263 | 97251.725460 | 40.399320 | 1.69e-009 | 1.000000 |
| SYM+I+G | 48496.12858 | 265 | 97522.257160 | 310.931020 | 3.03e-068 | 1.000000 |
| SYM+G | 48507.86910 | 264 | 97543.738200 | 332.412060 | 6.57e-073 | 1.000000 |
| K80+I+G | 48565.20726 | 261 | 97652.414520 | 441.088380 | 1.65e-096 | 1.000000 |
| K80+G | 48577.92428 | 260 | 97675.848560 | 464.522420 | 1.35e-101 | 1.000000 |
| F81+I+G | 48904.21932 | 263 | 98334.438640 | 1123.112500 | 0.00e+000 | 1.000000 |
| F81+G | 48920.44927 | 262 | 98364.898540 | 1153.572400 | 0.00e+000 | 1.000000 |

…

-lnL: negative log likelihod

K: number of estimated parameters

AIC: Akaike Information Criterion

delta: AIC difference

weight:AIC weight

cumWeight: cumulative AIC weight
